# Input-dependent directionality of interactions between cortical areas

**DOI:** 10.64898/2026.05.18.725829

**Authors:** Francesca Mastrogiuseppe, Joana Carmona, Byron M. Yu, Adam Kohn, Christian K. Machens

## Abstract

Studying the temporal coordination of activity signals across areas is essential for understanding brain function. Recent studies using cross-covariances show that activity directionality can shift rapidly with behavioral or task demands; yet, the circuit mechanisms underlying these changes remain unclear. Here, we use recurrent network models to investigate how directional interactions emerge and are flexibly reconfigured in multi-area cortical circuits. We show that, for fixed connectivity, directionality is shaped by how common inputs align with recurrent connectivity and the associated internal timescales of activity. In multi-area circuits with locally balanced excitation and inhibition, this reveals a predominant role for inputs to excitatory over inhibitory populations in controlling directionality. These inputs govern the directionality of the latent signals that account for most of the shared activity across areas, predominantly reflecting widespread and coherent activity fluctuations. Our models capture key features of cross-covariances from primate areas V1 and V2 and suggest parsimonious candidate mechanisms for the shift in directionality reported in these areas. This work establishes a mechanistic framework for understanding dynamic changes in temporal coordination across brain areas.

## Introduction

Explaining how the brain processes information to control behavior requires understanding how neurons across different areas interact. Time-resolved cross-covariance functions of neural activity, along with their normalized counterparts (cross-correlations), provide a simple yet powerful tool to assess these interactions. While the magnitude of a cross-covariance reflects the strength of statistical, and therefore functional, interactions between two recorded areas, its shape reflects the temporal ordering of these interactions. In particular, an asymmetric cross-covariance, with greater values for positive than for negative lags (or vice versa), indicates that fluctuations in one area tend to precede those in the other, thereby defining an emergent directionality in the temporal coordination of their activity.

Cross-covariance functions have long been central to the analysis of neural dynamics, both within local [1, 2, 3, 4, 5] and mesoscopic, multi-area cortical circuits [6, 7, 8, 9, 10]. Recently, interest in the latter application has been renewed [11, 12, 13, 14, 15], driven by the increasing availability of large-scale population recordings spanning multiple areas [16, 17, 18, 19, 20]. In this setting, cross-covariances are often computed from latent signals that summarize activity locally within each area [13, 21, 22, 15], reflecting for example the mean population activity [11, 23] or latent representations of sensory and cognitive variables [15, 20].

An important feature of inter-areal cross-covariances is their dynamic and context-dependent nature: their temporal structure and directionality can shift on short timescales depending on sensory inputs or task demands. For instance, van Kempen et al [11] found that cross-covariances between populations in primate V1 and V4 are nearly symmetric during fixation but become asymmetric – showing a net directional influence from V4 to V1 – during selective attention. Semedo et al [13] reported that directionality between areas V1 and V2 reverses between stimulus-driven and spontaneous activity, shifting from V1 leading V2 to V1 following V2. How these changes in functional interactions relate to the structure and operating regimes of the underlying cortical circuitry remains, however, largely unknown.

Experimentally measured cross-covariances are influenced not only by the connectivity scaffold across areas, which is largely stable over the timescale of a single trial, but also fast circuit-level processes [24, 25, 20]. Key contributors include inputs from other brain regions conveying sensory [4] as well as internal or cognitive signals [26, 25, 27, 28]. From both experimental and modeling perspectives, understanding how these inputs interact with the recurrent architecture of mesoscale cortical circuits to shape directionality remains a major challenge. This is because cortical areas are interconnected through bidirectional, recurrent synaptic pathways, such that any input reaching two areas simultaneously can reverberate repeatedly between them, and the resulting directionality reflects only the net outcome of these complex interactions.

In this work, we establish a modeling framework based on recurrent neural networks to elucidate how directional interactions, as measured by cross-covariances, emerge and can be dynamically reconfigured in multi-area cortical circuits. We first establish a general theory that provides both a quantitative and intuitive account of directional interactions and their dependence on common inputs. The theory shows that directionality can be tuned by varying how inputs align with network connectivity. We apply these insights to models of interconnected cortical areas with excitatory-inhibitory and feature-specific connectivity. Our analysis provides two key insights. First, in networks with balanced within-area connectivity, directionality of across-area interactions is primarily shaped by inputs targeting excitatory populations. Inputs to inhibitory populations control other activity features, such as amplitudes and intrinsic timescales [29, 30]. Second, this input-dependent directionality bias is robustly expressed by latent signals that are maximally correlated across areas, which dominate interactions at the population level. These signals encode widespread fluctuations in firing rate that coherently drive the activity of all neurons in the two areas up and down [31, 11, 32]. Using simultaneous recordings from primate areas V1 and V2 [9], we show that our models capture significant structure in the V1-V2 population activity, and allow us to derive candidate circuit mechanisms compatible with the shift in directionality between stimulus-driven and spontaneous activity reported in earlier work [13].

## Results

We consider experiments in which neural activity from two brain areas is recorded simultaneously over time (Fig. 1a). For each area, activity consists of continuous signals representing binned spikes from a single neuron or latent population signals (Fig. 1a-b); the latter encode patterns of co-variability within an area and are extracted from multiple neurons via dimensionality reduction [21, 13, 22]. Given activity signals *x*_*i*_(*t*) and *x*_*j*_(*t*) from two areas, the cross-covariance function (Fig. 1c) is defined as:

**Figure 1:**
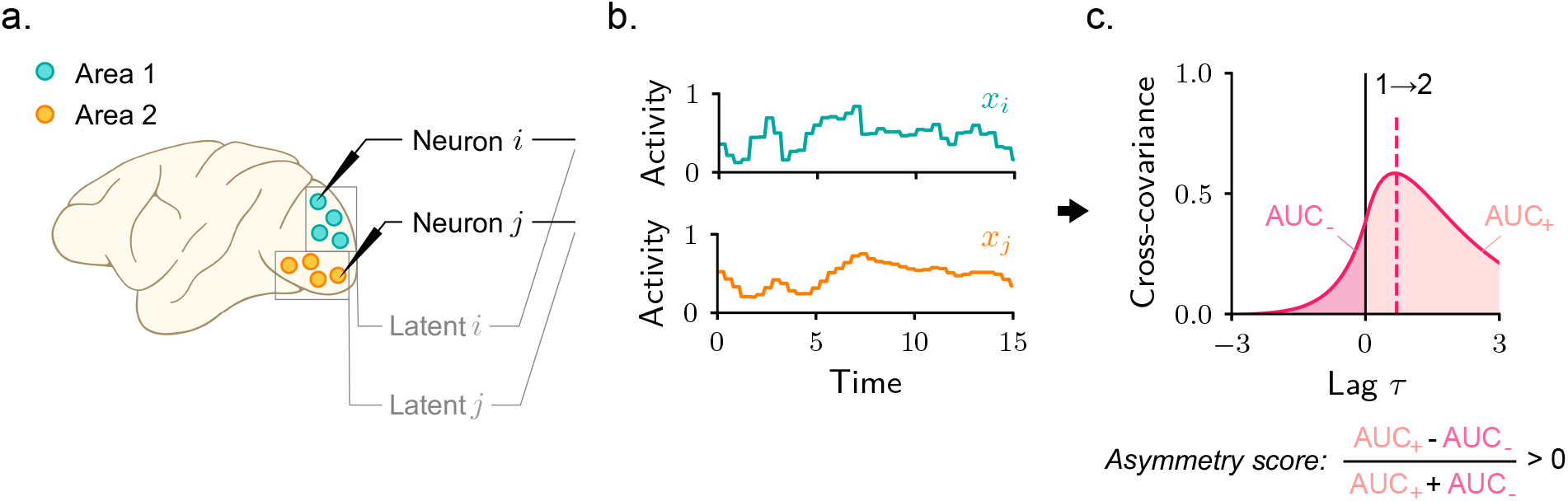
Estimating directionality of cortico-cortical interactions via cross-covariances. **a**–**b**. Simultaneous population recordings from pairs of cortical areas. Activity signals correspond to single neurons (black) or entire populations of neurons (grey) in the form of latent signals. **c**. Time-resolved cross-covariance between the two activity signals in b. Activity signals are aligned in time to the trial onset, and averages are computed over trials. We quantify directionality by computing an asymmetry score defined as the normalized difference between the areas under the cross-covariance curve for positive (AUC_+_) and negative (AUC_*−*_) lags. The value of this score is independent of how cross-covariances are normalized, and would therefore be identical if computed from cross-correlations instead.

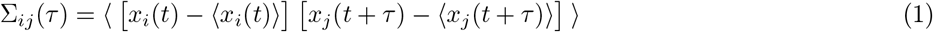

where *τ* is a time lag and angular brackets indicate averages over trials. Our focus is on the temporal structure of cross-covariances, particularly their symmetry around zero lag. An asymmetric (resp. symmetric) cross-covariance indicates that functional interactions are directional (resp. non-directional). We quantify asymmetry using a standard metric: the normalized difference between the area under the curve for positive and negative lags [33, 8, 13] (Fig. 1c). In our convention, a positive asymmetry score implies that the activity in area 1 tends to precede that of area 2, signaling an effective flow from the first area to the second one.

### Input-dependent directionality of interactions in recurrent circuits

Recent experimental findings have shown that the temporal structure of cross-covariances can undergo major qualitative changes when computed over trial epochs with different task or stimulus conditions [11, 13]. To identify the circuit basis of cross-covariance directionality in multi-area networks and the mechanisms that enable its rapid reconfiguration, we consider recurrent neural network models obeying [34, 35, 36]

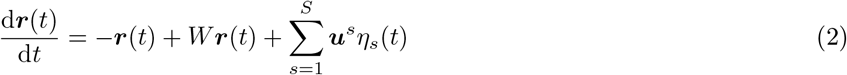

where ***r*** is a vector representing activity of units in the two areas, and *W* is a matrix encoding together within- and across-area synaptic connectivity, which we assume to be fixed (Fig. 2a, see Methods). The circuit receives inputs from external sources, modeled as time-varying, statistically identical and independent white noise signals *η*_*s*_(*t*). Each source influences activity via an input vector ***u***^*s*^, specifying which units are targeted and with what strength. These inputs drive time-varying, coordinated activity within the circuit, which we quantify using cross-covariances.

**Figure 2:**
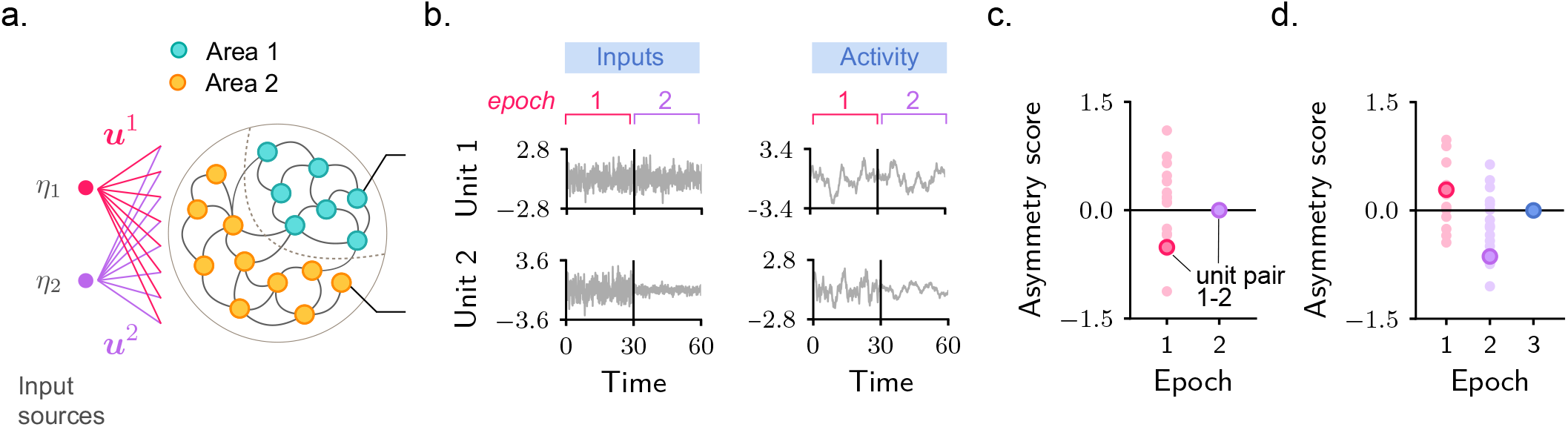
Recurrent neural network modeling setup. **a**. Schematic of a circuit in which units are grouped in two areas and receive input from two sources. **b**. Simulated external inputs (left) and output activity (right) for two units in an example circuit. Activity is simulated over two consecutive epochs, with only one input source active in each epoch. In the simulated circuit, each area consists of four units, and within- and across-area connectivity is drawn at random (see Methods). The input vectors for the two epochs are chosen to align differently with respect to connectivity (for details, see Fig. 3e and corresponding text). **c**. Cross-covariance asymmetry scores for pairs of units in different areas. Solid dot: pair of units 1 and 2 shown in b; transparent dots: remaining pairs. Most pairs exhibit strongly directional interactions in the first epoch, but not in the second. Cross-covariances are estimated by simulating activity over 100 trials, each with a distinct initial condition and input realization, and then applying Equation 1. **d**. Cross-covariance asymmetry scores after shuffling the recurrent connectivity while keeping the inputs fixed. Interactions are strongly directional in both the first and second epochs. Directionality can still be abolished for an appropriately chosen input vector (third epoch).

The directionality of activity within these models is, in general, hard to predict. Once entering the circuit, input signals do not follow a simple directed transmission path but instead reverberate repeatedly within and across areas through recurrent connections. To build intuition, we begin by examining an example circuit with randomly chosen connectivity (biologically constrained models are introduced in later sections). We simulate activity over many trials, each consisting of two distinct epochs. Each epoch is associated with a different input source and corresponding input vector – for example, one epoch may reflect a state-related input that engages units in both areas uniformly, whereas the other may reflect a sensory input that preferentially targets a subset of units in a single area. Input and activity traces for one representative unit in each area are shown in Fig. 2b. For each epoch, we quantify the directionality between these two units, as well as other across-area pairs, using the cross-covariance asymmetry score (Fig. 2c).

The two epochs exhibit markedly different directionality properties. Cross-covariances among most pairs of units show strong temporal asymmetry in the first epoch (Fig. 2c, magenta), but only weak asymmetry in the second (Fig. 2c, violet). Importantly, across the two epochs, synaptic connectivity is held fixed, and the input signals have similar temporal structure. The shift in directionality must therefore arise from the change in input vector – that is, from a change in which units are most strongly or weakly driven by the input source. More specifically, the different directionalities seen across epochs arise from a change in the alignment between the direction of the input vector and the fixed structure set by recurrent connectivity. To demonstrate this, we repeat the analysis in a model where inputs are held fixed but recurrent connectivity weights are shuffled. In this circuit, across-area cross-covariances display strong directionality in both epochs (Fig. 2d). Reduced directionality can however still occur for alternative input directions (Fig. 2d, blue).

These simulations suggest that external inputs can flexibly shape the directionality of interactions in multi-area circuits with fixed connectivity. This modulation induces qualitative changes even within the timescale of a single trial, consistent with experimental observations [11, 13]. To elucidate the mechanisms underlying these effects, we developed a theoretical framework that offers both an intuitive and quantitatively precise account of input-dependent directionality.

### Mechanisms underlying input dependence of directional and non-directional interactions

Consider the process through which a circuit as in Fig. 2 transforms inputs into output activity. Although each unit is directly influenced by the input, its activity reflects a transformed version of that signal, resulting from reverberation through recurrent connections (Fig. 3a). To examine this transformation in detail, we analyze activity in the transformed space defined by the eigenvectors of the matrix representing synaptic connectivity within and across areas (Fig. 3b). The key advantage of this space is that the mapping from inputs (left) to activity (right) becomes straightforward: for each eigenvector, the activity is obtained by filtering the input with a timescale set by the associated eigenvalue [37] (Fig. 3b, see Methods). The components of transformed activity therefore consist of input signals filtered at distinct timescales. In Fig. 3b, for example, the two eigenvectors are associated with a large and a small eigenvalue, giving rise to activity components with slow and fast temporal dynamics, respectively (see bottom panel for comparison).

**Figure 3:**
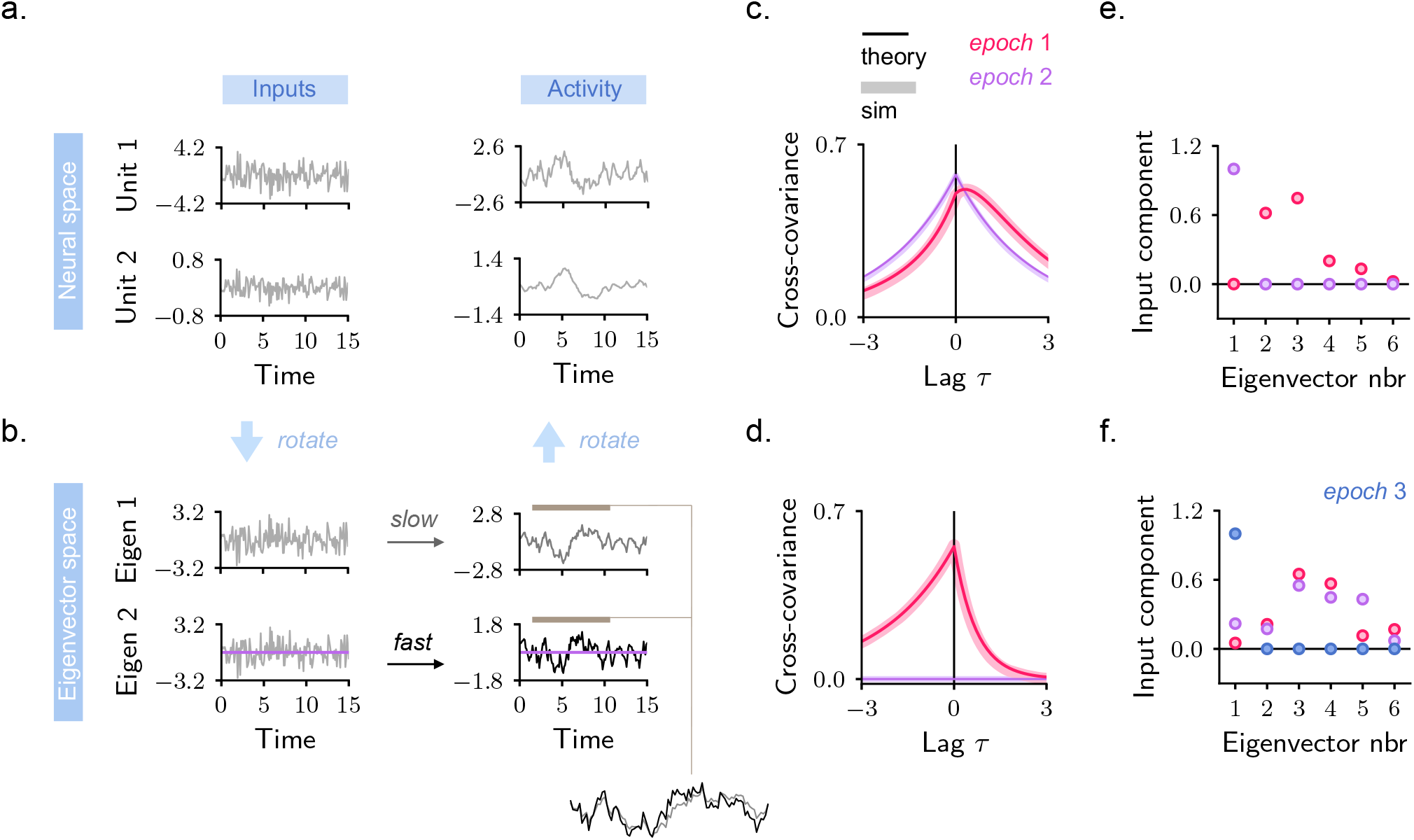
A theoretical framework for directionality and its dependence on external inputs. **a**. Inputs (left) and output activity (right) for two sample units from an example circuit model. **b**. Inputs and activity represented in the transformed space defined by the eigenvectors of the synaptic connectivity matrix. For clarity, only components associated with the first two eigenvectors are shown. In the transformed space, activity is obtained by linearly filtering the corresponding input signals, with a timescale determined by the eigenvalues (see Methods). Here, the first (resp. second) eigenvector is associated with a large (resp. small) eigenvalue, producing a slow (resp. fast) activity trace (see bottom right panel for comparison). In a and b, grey lines correspond to the first simulation epoch. Violet lines correspond to the second epoch; to reduce clutter, these are only shown in the bottom row of b. Note that, formally, the transformation from the neural to the eigenvector space is not always a rotation. This is because, in all biologically-relevant circuits considered here, connectivity matrices are non-normal and eigenvectors are not orthogonal. This also implies that the dimensions of activity involved in the transformation from the neural to the eigenvector space, and back, are not identical (see Methods). **c**. Cross-covariance between the two activity signals shown in a. Thick lines are computed from simulated activity; thin lines from theory. Lags are measured in arbitrary time units. **d**. Same as c, but for activity signals in the eigenvector space (shown in b). **e**. Components of the input used in Fig. 2 in the eigenvector space. **f**. Same as e, for shuffled connectivity (Fig. 2d).

This transformed representation of activity provides an intuitive explanation for the emergence of directional interactions. Indeed, activity components in the transformed space display systematic temporal relationships: components associated with slower eigenvectors tend to lag behind those linked to faster ones, resulting in asymmetric cross-covariances (Fig. 3d, magenta). When mapped back into neural space, these temporally structured components combine linearly to form the activity of individual neurons (Fig. 3a). Because the combination generally differs across neurons – and in particular across neurons in different areas – lead-lag relationships and asymmetric cross-covariances appear in the neural space as well (Fig. 3c, magenta), with units that more strongly express fast components leading the interactions.

Our analysis highlights how recurrent circuits process inputs at multiple timescales, providing a substrate for directional interactions. The same framework also explains when directional interactions arise in practice (e.g., first simulation epoch in Fig. 2b) and when they do not (second epoch). During the first epoch, activity in the eigenvector space behaves as previously described: each eigenvector processes the input at a distinct timescale (Fig. 3b, grey and black), giving rise to a temporal hierarchy that manifests as directional interactions in the transformed and neural spaces (Fig. 3c–d, magenta). In the second epoch, by contrast, only the first eigenvector is associated with an input signal, while inputs to the remaining eigenvectors vanish (Fig. 3b, violet). Consequently, the circuit expresses a single filtered version of the input, and a single timescale. When mapped back into neural space, this results in synchronized activity and non-directional cross-covariances (Fig. 3c, violet). Importantly, the two epochs correspond to different choices of the input vector: during the first epoch, the input vector was chosen to have strong components along many eigenvectors (Fig. 3e, magenta), whereas in the second it aligns primarily with the first eigenvector (Fig. 3e, violet). When recurrent connectivity is shuffled (Fig. 2d) the eigenvector directions are modified. In this case, the input vectors acquire components along multiple eigenvectors in both epochs (Fig. 3f), leading to directional interactions (Fig. 2d). By contrast, aligning the input with an eigenvector of the shuffled connectivity (Fig. 3f, blue) yields again non-directional interactions (Fig. 2d, blue).

We conclude that inputs aligned with single eigenvectors of the connectivity matrix promote non-directional interactions. Although the scenario considered so far involved a single input source active per epoch, similar results apply in the presence of multiple input sources (Supplementary Fig. 2). More generally, our theory indicates that the alignment between the input and eigenvector directions is a key determinant of directionality, as it sets the number of effective timescales and, consequently, the temporal complexity that supports directional interactions. Controlling the alignment between inputs and eigenvectors represents therefore a mechanism for flexible and rapid reconfigurations of directionality.

### Inputs to excitatory populations control across-area directionality in locally balanced circuits

Our findings so far suggest a simple strategy for understanding directional interactions in recurrent circuits: characterize the structure of the connectivity eigenvectors, determine how strongly each eigenvector is recruited by the external inputs, and reconstruct activity in neural space by combining the contributions of the different eigenvectors. In the following, we apply this framework to circuits composed of two cortical areas. For the simplified models considered here, the eigenvector structure can be characterized explicitly. Individual eigenvectors are not associated with specific areas, but rather involve units distributed across both areas [38]. Consequently, the associated timescales are also spatially distributed rather than localized. The details of this analysis are provided in the Methods; here, we discuss the main results.

We begin with a circuit of two areas, each containing two units, representing one excitatory (E) and one inhibitory (I) population (Fig. 4a). Within each area, recurrent interactions are strong, with excitatory connections counteracting inhibitory ones [39, 34]. Inter-areal connectivity is instead purely excitatory [32]. Because excitatory neurons constitute the majority of cortical cells and are most commonly recorded in experiments, we focus on the excitatory populations in each area and quantify directionality through their cross-covariance. Input vectors to this circuit are four-dimensional, with each entry quantifying the input strength onto the two excitatory and two inhibitory populations. In the cortex, afferent inputs target both excitatory and inhibitory populations within recipient areas [40] – a pattern thought to stabilize activity amplitude and integration timescales across the cortical hierarchy [29, 30, 41]. Here we test whether, and in what way, inputs to excitatory and inhibitory populations influence across-area directionality. To this end, we systematically vary the strength of inputs to excitatory and inhibitory populations in the two areas. These manipulations modify the alignment of the input vector with respect to the eigenvectors of the synaptic connectivity, which our theory predicts can in turn affect directionality.

**Figure 4:**
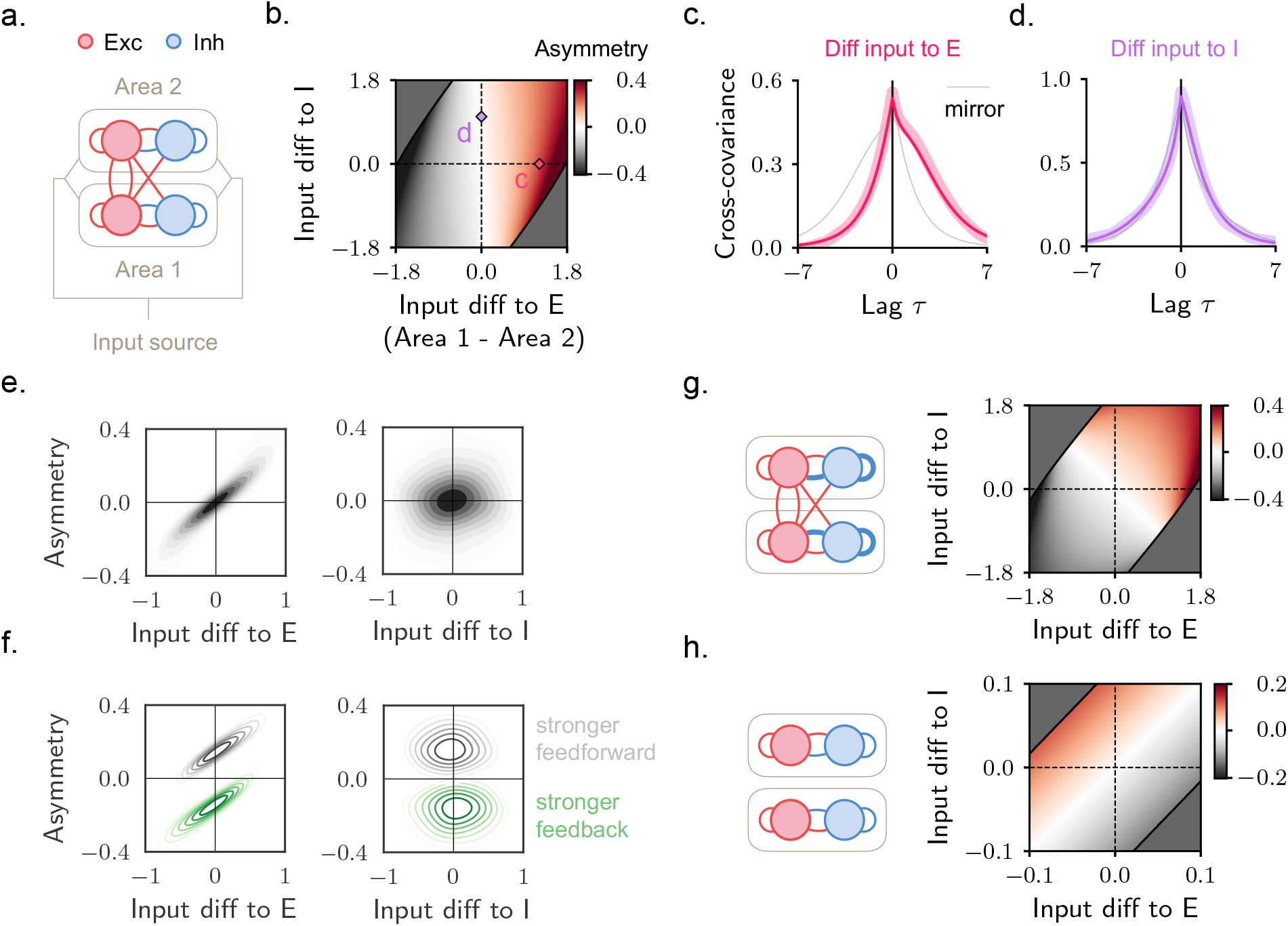
Across-area directionality in two-area excitatory-inhibitory circuits. **a**. Schematic of the circuit with a common input signal. The across-area cross-covariance is measured from the activity of the two excitatory populations. **b**. Cross-covariance asymmetry score as a function of the difference in input to excitatory (horizontal axis) and inhibitory (vertical axis) populations. We start from a configuration where the input strength is equal to 1 for all populations, and then gradually increase the input to a population in one area while equally decreasing it for the corresponding population in the other area. (A complementary approach and quantification is provided in Supplementary Fig. 5). The dark grey region corresponds to a regime in which excessively strong inhibitory inputs suppress activity, yielding negative correlations between excitatory and inhibitory activity. As this correlation is typically positive in cortical circuits, we exclude this region from the analysis (see Methods for details). **c**. Cross-covariance for unequal input to excitatory populations and equal input to inhibitory ones (magenta diamond in panel b). Grey line represents the mirror image of the magenta one; the mismatch between the two provides an intuitive visualization of asymmetry. **d**. Same as c, but for unequal input to inhibitory populations and equal input to excitatory ones (violet diamond in panel b). **e**. Asymmetry score as a function of the difference in input strength to excitatory (left) or inhibitory (right) populations. Density plot for many input configurations, sampled uniformly at random. **f**. Same as e, but for two different circuits, in which feedforward (grey) or feedback (green) connectivity is stronger. **g**. Same as in b, but for a circuit with a stronger dominance of inhibitory over excitatory connections (circuit schematic on the left). **h**. Same as in b, but for a circuit with balanced within-area connectivity and no inter-areal connections.

We begin from a baseline configuration in which all excitatory and inhibitory populations receive identical input (Fig. 4b). We then gradually increase the input to the excitatory population in one area while decreasing it in the other (difference shown on horizontal axis), and apply an analogous manipulation to inhibitory populations (vertical axis). Differences in input to excitatory populations have a pronounced impact on directionality: when the input difference is large, interactions are always strongly directional, with the most strongly driven area leading in time. In contrast, differences in input to inhibitory populations only exert a minor influence: even when this difference is large, directionality remains primarily determined by excitatory inputs. Example cross-covariances in which the input difference affects only excitatory or inhibitory populations are shown in Fig. 4c–d, respectively. Consistent with Fig. 4b, a pronounced asymmetry is observed in C, but not in D. Thus, directionality exhibits unequal sensitivity to inputs targeting excitatory versus inhibitory populations.

As a complementary quantification, we consider many randomly generated inputs and compute the difference in strength between the two excitatory (Fig. 4e, left) or inhibitory (Fig. 4e, right) populations, together with the corresponding asymmetry score. Large differences in inputs to the excitatory populations are consistently associated with strong directional biases, directed from the area receiving the largest input to the other (Fig. 4e, left). In contrast, differences in inputs to inhibitory populations show no clear monotonic relationship with directionality (Fig. 4e, right). These results are obtained for a circuit with symmetric inter-area connectivity. In models where the feedforward (Fig. 4f, grey) or the feedback (Fig. 4f, green) pathway is stronger, interactions are biased so that the area from which the strongest connectivity originates leads in time. Nonetheless, in these models too, differences in inputs to the two excitatory populations bias directionality to be led by area receiving stronger input, and can even reverse the directionality imposed by the connectivity. Differences in inputs to the two inhibitory populations produce instead no additional bias (for a systematic parameter analysis, see Supplementary Fig. 6).

Intuitively, one might expect directionality to be less sensitive to inputs targeting inhibitory populations because these populations do not project across areas, and thus influence inter-areal dynamics only indirectly. This intuition holds when inter-areal connections are much stronger than within-area ones (see Methods), but does not hold in the circuit considered here, where strong local coupling allows inhibitory inputs to influence excitatory activity and inter-areal interactions. To demonstrate this, we repeat the analysis from Fig. 4b in a circuit where the local excitation-inhibition balance (i.e. the matching in strength of within-area excitatory and inhibition connections) is less precise, and activity is more strongly inhibition-dominated (Fig. 4g). In this case, the asymmetry between excitatory and inhibitory inputs is markedly reduced: inputs to either population modulate directionality, biasing the area receiving the stronger input to lead (Fig. 4g). Thus, excitatory inter-areal connections alone cannot explain the unequal sensitivity; local excitation-inhibition balance is also required. Nevertheless, excitatory inter-areal connectivity is a key ingredient. To illustrate this, we consider a circuit in which within-area balance is restored but inter-areal connections are removed, so that across-area covariability arises solely from shared inputs (Fig. 4h). In this case, directionality is again equally sensitive to excitatory and inhibitory inputs (this time, with the two types of input influencing directionality in opposite directions, see Methods). Together, these results indicate that the dominant influence of inputs to excitatory populations emerges specifically in circuits that combine locally balanced dynamics and excitatory inter-areal projections.

Our mathematical framework provides a useful tool for understanding these results. In Fig. 5a (resp. b), we visualize the activity underlying the cross-covariances in Fig. 4c (resp. d), where inputs differentially target excitatory (resp. inhibitory) populations. The activity traces (top) can be decomposed into two eigenvector signals (bottom), each associated with a distinct timescale. Our theory indicates that directional interactions emerge when activity spans multiple eigenvectors and timescales, whereas the dominance of a single eigenvector results in non-directional interactions. Consistent with this, when excitatory populations receive distinct inputs (Fig. 5a), both eigenvector components are strongly expressed, producing an asymmetric cross-covariance (Fig. 4c). By contrast, when only inhibitory inputs differ (Fig. 5b), activity is dominated by a single eigenvector signal, yielding an approximately symmetric cross-covariance (Fig. 4d). Thus, differences in inputs to excitatory – but not inhibitory – populations give rise to multiple timescales, whose coexistence underlies directional interactions, with the area that more strongly expresses fast components leading in time.

**Figure 5:**
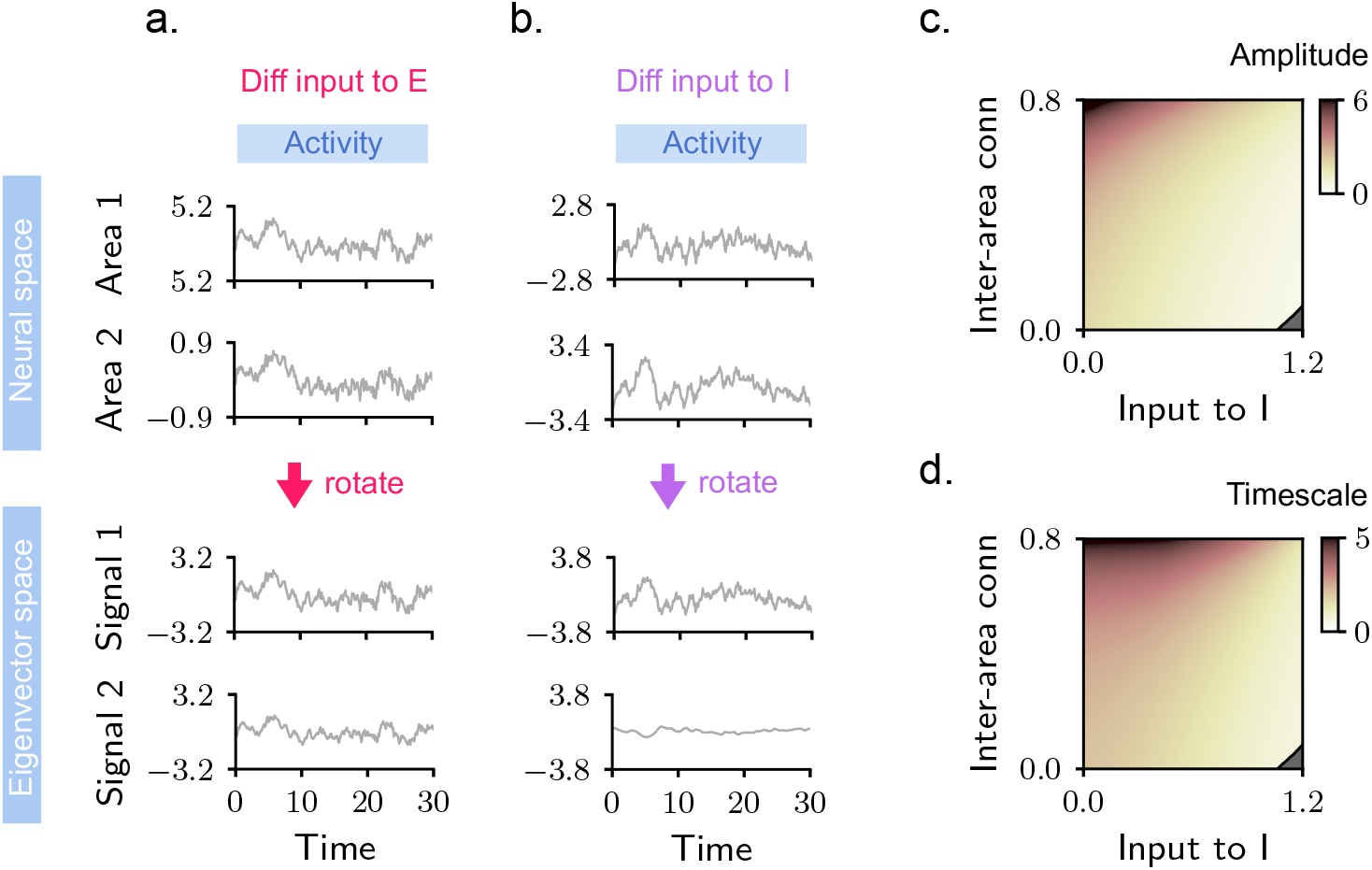
Directionality in circuits with balanced within-area connectivity. **a**–**b**. Simulated activity of the two excitatory populations for the input configurations shown in Fig. 4c–d. Top (resp. bottom): neural (resp. eigenvector) space. The amplitude and temporal structure of the second eigenvector signal in b is suppressed; a detailed analysis of this behaviour is reported in Methods. **c**. Activity amplitude as a function of the input to inhibitory populations (horizontal axis) and inter-areal connectivity strength (vertical axis). The input to the excitatory populations is held fixed. We show results for one of the two areas (the two areas yield qualitatively identical results). **d**. Same as c, but for the characteristic timescale of activity, computed as the width at half maximum of the auto-covariance.

We finally note that, although inputs to inhibitory populations exert only a limited influence on directionality, they nonetheless play an important role in mesoscopic dynamics (Fig. 5c–d). In particular, inhibitory inputs strongly modulate both the amplitude and the range of timescales of neural activity: increasing the inhibitory input (horizontal axis) leads to a reduction in both measures. This modulation counteracts the tendency of amplitude and timescale to grow in networks with increasing excitatory inter-areal connectivity (vertical axis) [32]. Thus, while not directly shaping directionality, inputs to inhibitory populations serve a key stabilizing role in mesoscopic cortical dynamics [40, 29, 30, 41].

### Feature-unspecific signals dominate across-area interactions and express input-dependent directionality

Our analysis so far indicates that activity directionality in locally balanced multi-area circuits is shaped by the strength of common inputs to excitatory populations. Unlike in the simplified model consider so far, excitatory populations in the cortex exhibit substantial structure: neurons are tuned to distinct features (e.g., different aspects of sensory stimuli), and both within- and across-area connectivity reflect this specificity [42, 43, 44, 45, 46, 47]. To examine how feature specificity influences directionality, we next consider multi-area circuits with feature-tuned populations. Fig. 6a illustrates an example with two features, A and B [32]. Each area includes one excitatory and one inhibitory population per feature. Both within and across areas, connectivity is stronger between populations tuned to the same feature. Within-area connectivity is balanced, while across-area connectivity is excitatory.

**Figure 6:**
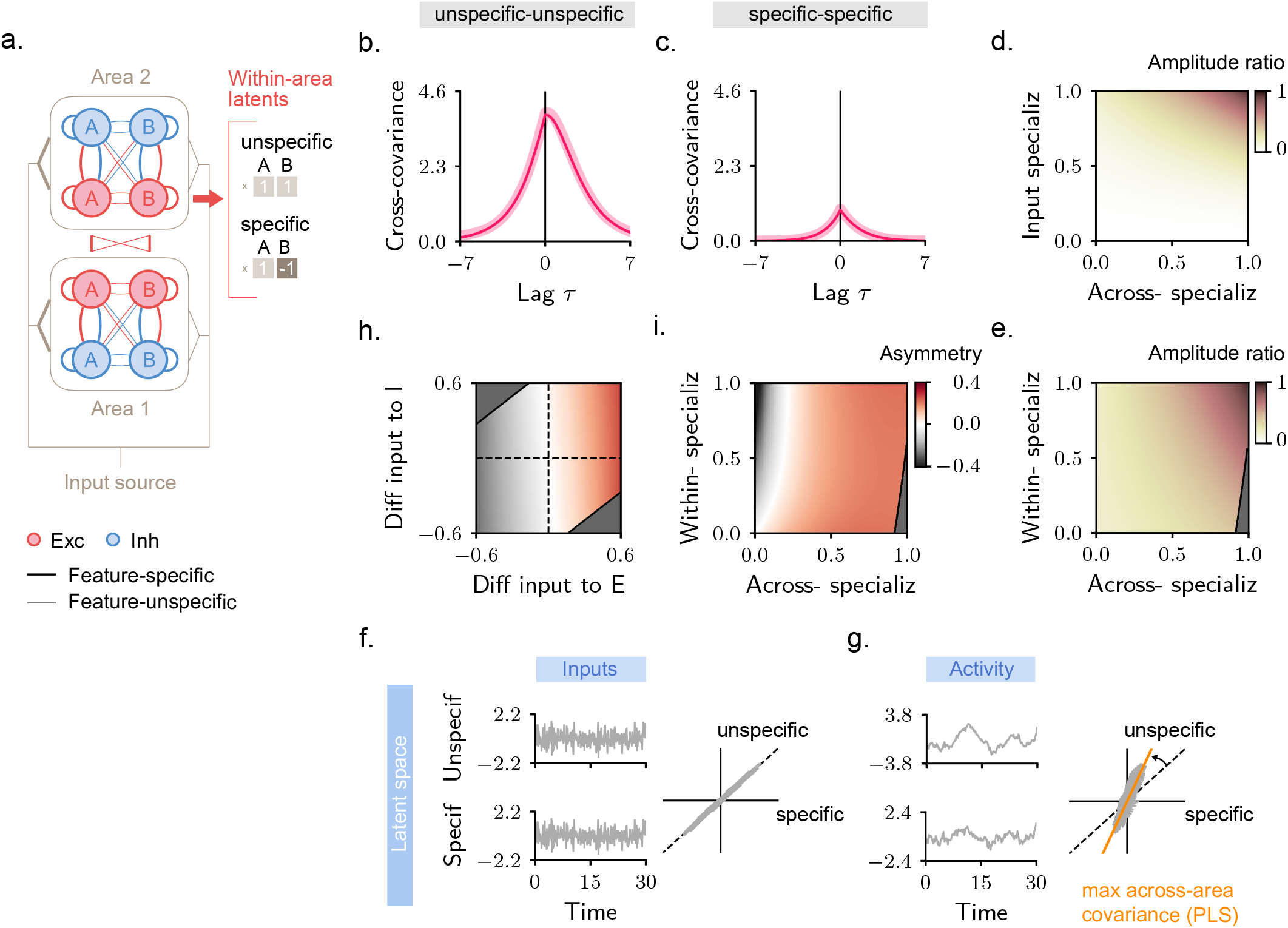
Across-area directionality in feature-specialized circuits. **a**. Left: schematic of a two-area circuit in which inputs and connectivity are tuned to features A and B. Across-area connectivity is shown schematically; it originates from all excitatory units and targets both excitatory and inhibitory populations in the other area, according to their feature tuning. The degree of feature specificity of inputs and connectivity is set by tunable parameters (see Methods), with 0 (resp. 1) corresponding to no (resp. full) feature specificity. These are varied in panels d, e, i. Right: the activity of the two excitatory populations in each area is described in terms of a feature-unspecific and a feature-specific latent. **b**–**c**. Example cross-covariance computed from feature-unspecific (b) and feature-specific (c) latents. **d**–**e**. Ratio between the amplitude of feature-specific and unspecific cross-covariances (taken as their maximum values) as a function of the degree of feature specialization in across-area connectivity and inputs (d), or in within- and across-area connectivity (e). In all cases, the ratio is bounded between 0 and 1. **f**–**g**. Simulated inputs (f) and activity (g) for excitatory populations in one of the two areas. Insets on the right show a time-implicit view of the temporal traces depicted on the left. The orange line denotes the axis identified by the Partial Least Squares (PLS) algorithm, trained to identify the directions of maximal covariance between areas. **h**. Asymmetry score of the cross-covariance from feature-unspecific latents as a function of the difference in overall input to excitatory and inhibitory populations. Note the similarity with Fig. 4b. These results do not qualitatively depend on the degree of specialization of within- and across-area connectivity (see Methods). **i**. Asymmetry score of the cross-covariance computed from feature-specific latents as a function of the degree of specialization of within- and across-area connectivity. We considered a fixed input configuration as in b–c, for which inputs preferentially target A over B populations, and the overall input to excitatory populations is larger for area 1 than area 2, implying positive asymmetry for unspecific latents (h).

To link these models with recent work addressing across-area interactions at the population level [11, 13, 15, 48, 20], we characterize activity using latent variables (Fig. 6a, right). Specifically, for each area, we define a feature-unspecific latent by projecting activity of the two A and B excitatory populations onto the axis [1, 1]. This latent – equivalent to the mean activity of excitatory neurons within one area – encodes fluctuations that modulate coherently the activity of differently tuned populations. We also define a feature-specific latent along the axis [1, −1], capturing fluctuations that modulate the activity of differently tuned populations in opposite directions. Across-area interactions are then quantified via cross-covariances between pairs of unspecific (Fig. 6b) and specific latents (Fig. 6c).

We begin by assessing how much these two sets of latents contribute to across-area interactions by comparing the amplitude of their cross-covariances (Fig. 6b–c). Across models with varying degrees of connectivity and input specialization, the ratio between specific and unspecific amplitudes is consistently below one (Fig. 6d–e). This indicates that across-area interactions are dominated by feature-independent signals. To clarify the origin of this imbalanced contribution, we visualize how the circuit transforms inputs into output activity. Focusing on one area, we represent inputs (Fig. 6f) and output activity (Fig. 6g) in terms of their feature-unspecific and specific latent components. In the simulated circuit, these components have comparable amplitudes at the input stage, but not in the output activity, where the unspecific component becomes dominant. Recurrent interactions therefore selectively amplify feature-unspecific fluctuations, causing most of the variance within each area to be captured by unspecific signals, as recently demonstrated in Javadzadeh et al [32]. Because these signals are coordinated across areas, they also account for most of the variance that is shared between areas, resulting in large-amplitude cross-covariances (Fig. 6d–e). In Fig. 6g, we show the activity direction identified by a dimensionality reduction algorithm, Partial Least Squares (PLS), trained to maximize covariance across areas [13, 19]. Consistent with our analysis, this direction aligns more strongly with the unspecific than the specific direction.

We next examine the directionality of feature-unspecific and feature-specific latents. The behavior of feature-unspecific latents (Fig. 6h) closely mirrors that seen in simpler, feature-independent circuits (Fig. 4b). In particular, directionality is strongly influenced by inputs to excitatory populations, with the area receiving the larger overall input leading in time, while inputs to inhibitory populations have only a weak effect. This result is independent of the circuit’s degree of feature specialization and holds for any form of feature-specific inputs and connectivity (see Methods and Supplementary Fig. 8). Thus, even in structured circuits, inputs to excitatory populations remain a key determinant of directionality, shaping cross-covariances of the dominant, feature-unspecific activity signals.

In contrast, the directionality of feature-specific latents depends not only on inputs but also on the fine feature-dependent structure of the circuit. Consider, for example, the input configurations resulting in the cross-covariances shown in Fig. 6b–c, in which excitatory populations in the first area receive stronger input, and therefore lead the feature-unspecific cross-covariance. For this configuration, the feature-specific cross-covariance can display either directionality depending on the degree of feature specificity within and across areas (Fig. 6i). In particular, feature-unspecific and specific latents share the same directionality only when across-area connections are sufficiently feature-specific relative to within-area ones (Fig. 6i, red region), whereas weakly feature-specific connectivity can reverse the directionality of feature-specific latents relative to unspecific ones (Fig. 6i, black region).

In summary, in structured multi-area circuits, directionality remains primarily shaped by differences in input strength to excitatory populations. These inputs set the directionality of the dominant cross-covariance, corresponding to feature-unspecific signals. The feature-specific structure of the circuit also influences directionality. This effect is, however, only observed in the cross-covariance of feature-specific signals, which have lower amplitude.

### Models capture cross-covariance structure and suggest plausible mechanisms for V1–V2 recordings

We set out to compare our models to data from simultaneous population recordings in areas V1 and V2 of anesthetized primates [9, 13] (Fig. 7a). In these recordings, each trial includes a stimulus epoch, during which a drifting sinusoidal grating is presented, followed by an epoch without stimulation (Fig. 7b). Recent work showed that the dominant across-area cross-covariance, identified via canonical correlation analysis [49], displays a temporal asymmetry primarily directed from V1 to V2 soon after stimulus onset, and from V2 to V1 during the spontaneous epoch [13]. Our goal is to employ our theoretical framework to link these empirical observations to circuit mechanisms.

**Figure 7:**
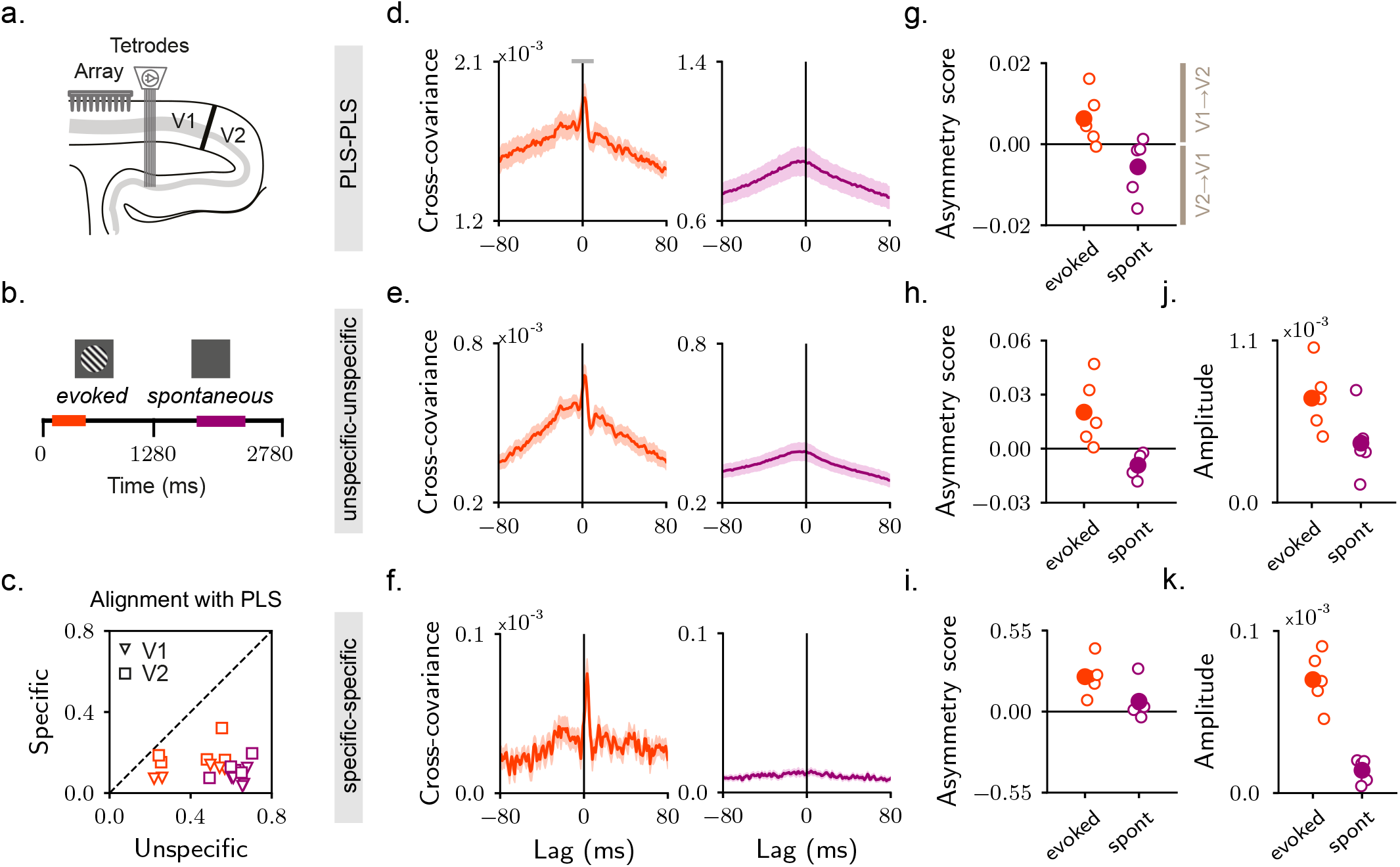
Validating the model on simultaneous population recordings from primate areas V1 and V2. **a**. Schematic of experimental setup [9]. Activity is recorded simultaneously in V1 and V2 from anesthetized primates. We summarize activity through latent signals, obtained by projecting activity within each area along two directions: a feature-unspecific and a feature-specific one. Details on how these directions are computed are provided in Methods. We also use a dimensionality-reduction algorithm, Partial Least Squares (PLS), to identify the latent signals within each area that account for the majority of shared variance across areas. **b**. Each trial includes a period of visual stimulation (drifting sinusoidal gratings), followed by spontaneous activity. When comparing stimulus-evoked and spontaneous periods, we consider time intervals in the two epochs during which activity is almost stationary (colored bars in the timeline). Following Semedo et al [13], our stimulus interval corresponds to early evoked activity (late evoked activity yields more modest temporal asymmetries). Data include 5 sessions, each with 8 independent repetitions (different stimulus orientations). **c**. Alignment (computed as dot product between normalized vectors) between the PLS and feature-specific directions, compared with the overlap between the PLS and feature-unspecific directions. Triangles and squares represent latent signals in V1 and V2, respectively. Each symbol corresponds to one session (average over orientations). **d**–**f**. V1–V2 cross-covariances computed from the PLS (d), feature-unspecific (e), and feature-specific (f) latents. Solid lines represent the average cross-covariance across all datasets; shaded areas the s.e.m. across datasets. The grey horizontal bar in d indicates the lag window used to measure asymmetry scores. To capture inter-areal interactions at fast timescales, which dominate directionality in the stimulus epoch [13], we use a small window. **g**–**i**. Asymmetry scores of the cross-covariances computed from PLS (g), feature-unspecific (h), and feature-specific (i) latents. Each empty dot corresponds to one of the 5 session (average over 8 orientations). **j**–**k**. Amplitudes of the cross-covariances computed from feature-unspecific (j) and specific (k) latents, measured as their maximum value.

To begin, we validate our framework by showing that the V1–V2 population structure of recorded activity aligns with the models. We use PLS to identify activity directions that maximize across-area covariability. We also estimate feature-unspecific and feature-specific directions by assigning equal or opposite weight to units that are tuned to different features (A and B correspond here to the presented grating orientation and its orthogonal counterpart). The model predicts that the dominant PLS direction should align more strongly with the feature-unspecific than the feature-specific one (Fig. 6g). This prediction is borne out: the difference in alignment is consistent across areas and epochs, and strongest during the spontaneous epoch (Fig. 7c). Accordingly, the dominant PLS cross-covariance (Fig. 7d), previously characterized in Semedo et al [13], more closely resembles that computed from feature-unspecific latents (Fig. 7e) than feature-specific ones (Fig. 7f). In particular, the sign of its asymmetry score (Fig. 7g) is qualitatively consistent in both epochs with that of feature-unspecific (Fig. 7h) but not feature-specific latents (Fig. 7i). Overall, this analysis indicates that the dominant mode of V1–V2 interactions in the data reflects feature-agnostic fluctuations that jointly recruit differently tuned units across areas.

What circuit configuration can account for the empirical cross-covariances in Fig. 7d–f? To address this question, we closely examine cross-covariances during the two trial epochs and use model results from Fig. 6 to constrain the underlying circuitry. We begin by analyzing cross-covariance amplitudes. While the amplitude of the feature-unspecific cross-covariance is large and remains comparable across stimulus and spontaneous periods (Fig. 7j), the feature-specific one shows a marked decrease (Fig. 7k), becoming nearly negligible during the spontaneous epoch. Our model indicates that, for fixed connectivity, the amplitudes of the two cross-covariances can vary as a function of input specialization. In particular, cross-covariances from feature-specific latents can become vanishingly small if the input specialization is reduced (Fig. 6d). Overall, these observations are consistent with a model in which the dominant input to V1 and V2 is more strongly feature-specific during stimulus presentation than during spontaneous activity. Second, we analyze cross-covariance directionality. The feature-unspecific cross-covariance shows a systematic shift, from V1-leading early in the stimulus period (Fig. 7h, orange) to V2-leading in the spontaneous period (Fig. 7h, purple). In our model, the directionality of feature-unspecific latents is set by the input strengths to excitatory populations, with the area receiving the strongest input leading the interaction (Fig. 6h). Thus, the observed change in V1–V2 directionality is consistent with an input source whose preferential targeting of V1 over V2 is stronger during the stimulus than the sponteaneous period. Finally, during the stimulus period, the feature-specific cross-covariance also exhibits V1-leading directionality (Fig. 7i, orange). In the model, the directionality of these cross-covariances depends on the feature-specific structure of across-area connectivity (Fig. 6i). Specifically, the observed alignment in directionality between feature-unspecific and specific latents can be reproduced in a circuit where across-area connectivity is sufficiently feature-specific. Notably, recent experimental work provides evidence for such specificity in cortical networks [47], and theoretical work on activity timescales in the rodent visual cortex reached a similar conclusion [32].

Altogether, these considerations suggest a candidate mechanism compatible with the shift in directionality reported by Semedo et al [13] based on a change in the dominant input sources across the two epochs combined with feature-dependent structure of recurrent connectivity within and across areas. This mechanism is qualitatively consistent with the data, and broadly compatible with known anatomical organization [47, 50].

## Discussion

Recent advances in large-scale recording techniques [51], together with dimensionality reduction [21, 22, 52], now make it possible to estimate the inter-areal activity flow with high resolution, while relating these signals to internal and computational processes [15, 20]. Despite the progress, the link between the statistical estimates of this flow – typically based on time-lagged covariances – and the synaptic signaling occurring across areas at the circuit level has remained poorly understood. In this work, we propose a modeling framework based on recurrent neural networks for studying mechanistically the directional propagation of activity across areas.

We show that recurrent connectivity naturally promotes the emergence of directional interactions by supporting input processing at multiple timescales, which in turn gives rise to lead-lag relationships (Fig. 3a–d). Non-directional interactions emerge instead when a single timescale dominates. A key determinant of both the number of expressed timescales and directionality is the alignment between the input vectors and the eigenvectors of the synaptic connectivity matrix (Fig. 3e). Estimating eigenvectors requires precise knowledge of the synaptic structure and represents therefore a challenge for biological circuits. However, for simple structures such as the two-area excitatory-inhibitory networks considered here, eigenvectors can be computed explicitly (see Methods), providing insight into the input dependence of directionality. One important aspect of these eigenvectors is that they are not localized to single areas [38]: they have non-vanishing components across excitatory and inhibitory populations in both areas, implying that the associated timescales are expressed globally. At the same time, their spatial structure allows different timescales to be expressed with different strengths across areas, forming the substrate for directional interactions. Which timescales are ultimately expressed depends on how the external inputs recruit the different eigenvector signals. In particular, input configurations that differentially drive the excitatory populations of the two areas lead to different average timescales across areas and directional interactions from the faster area toward the slower one (Fig. 4c). Variability in average intrinsic timescales across cortical areas has been established experimentally [53, 54]. Our work connects timescales to activity propagation, and highlights their input-dependent and flexible nature.

Our analysis reveals that the excitatory-inhibitory structure of mesoscopic cortical circuits has precise implications for across-area directionality. In circuits with locally balanced E-I connectivity, only differences in inputs to excitatory populations affect directionality, whereas even large differences in inhibitory inputs fail to bias it. This is not because inputs to inhibitory populations are ineffective at shaping activity – they strongly influence its amplitude and timescale across areas (Fig. 5c–d, see also [55]). Rather, their alignment with respect to the connectivity eigenvectors leads to a single dominant timescale, entraining excitatory populations across areas (Fig. 5b). This mechanism relies on across-area connectivity that is excitatory and sufficiently strong, and is disrupted in non-balanced connectivities – highlighting how impairments to the local balance [56] can also modify across-area flow. Another consequence of the excitatory nature of across-area connectivity is that the dominant across-area cross-covariance (i.e. the cross-covariance computed from signals of maximal across-area covariability [13, 19]) predominantly reflects feature-unspecific latents, and therefore encodes fluctuations in the mean activity that are globally coordinated across areas (Fig. 6b–e). The preferential amplification of feature-unspecific latents through across-area excitatory connections (Fig. 6f–g) has been recently demonstrated in Javadzadeh et al [32]. Here, we show that the directionality of these latents depends on the strength of inputs to excitatory populations. In our models, across-area connectivity follows a simple like-to-like organization in both feedforward and feedback directions, consistent with experimental evidence [47]. However, several studies suggest that this connectivity is not exclusively like-to-like, but can also include preferential connections between neurons tuned to different features [57, 45, 58]. An important direction for future work is therefore to assess whether the dominance of feature-unspecific latents persists under more heterogeneous connectivity structures.

These results offer important insights on how directionality measures should be interpreted in the context of across-area interactions. In our models, directional interactions indicate that two areas express a shared signal at different average timescales (Fig. 3a–d). This differs from the classical “signal transmission” view, which assumes that a signal originates in one area and is then transmitted synaptically to another, where it appears after some delay. This perspective has recently informed the design of statistical models [21, 59]. Biophysically, it has a clear basis at millisecond timescales, but fails to capture the reverberation of signals through recurrent connections. Our framework aligns with this transmission perspective under specific conditions. For example, when the input targets only one area, that area contains both the fast input component along with a slower, recurrently processed one, whereas the other area expresses only the slower component and thus appears to lag (see Methods). Qualitatively, this mechanism strongly resembles feedforward transmission (although the cross-covariance displays smooth asymmetries rather than a sharp peak at short delays, see e.g. Fig. 4c). However, the two perspectives do not always coincide. For example, directionality fails to capture synaptic signaling via inhibitory units when within-area connectivity is balanced (Fig. 4b). Conversely, it can be non-zero even in the absence of inter-areal connectivity and transmission, provided that local timescales differ across areas (Fig. 4h). Directionality therefore reflects only specific modes of across-area signaling – a limitation that highlights the inherently functional nature of cross-covariance-based measurements [24]. Nonetheless, when interpreted in conjunction with theoretical models, these measures represent a potentially valuable tool for investigating mechanisms. In the V1–V2 data analyzed here, for example, directionality could be used to infer the feature specificity of inputs across epochs and to constrain the structure of inter-area connectivity (Fig. 7).

Our results provide insights for the design of experiments and the analysis of neural recordings. At the circuit level, our analysis yields experimental predictions about how input manipulations, such as optogenetic perturbations, should affect across-area directionality. First, perturbations targeting excitatory neurons should affect inter-area directionality more strongly than perturbations targeting inhibitory neurons. Second, the area receiving the strongest input should lead interactions, setting the directionality of the dominant, feature-unspecific cross-covariance. Third, the directionality of the feature-specific, lower-variance cross-covariance should only be affected by perturbations that are themselves feature-specific [60]. Crucially, these predictions are expected to hold only for pairs of areas that share strong bidirectional connectivity (Fig. 4h).

Our work has also implications for the use of latent variable models on multi-area data – an approach that is becoming increasingly common [13, 15, 19]. First, the directionality of a given latent can vary, even in a fixed recurrent circuit, due to changes in external inputs. For methods in which each latent is assigned fixed directionality [21], this implies that care must be taken in selecting analysis windows, which should correspond to periods of approximately stationary across-area signaling. Second, the variance associated with each latent is expected to depend on its loading structure. In particular, the model predicts that feature-unspecific latents, corresponding to single-neuron loadings with largely consistent signs, account for the largest fraction of shared variance within and across areas. Therefore, these latents are expected to be more readily reconstructed from neural recordings. While global, feature-unspecific fluctuations may be prominent [61], we emphasize the importance of studying lower-variance components, as they provide unique access to computationally relevant structure in both inputs and connectivity (Fig. 7).

Our analysis relies on a few key assumptions. First, our models do not incorporate explicit transmission delays arising from axonal and synaptic conduction. Even in the absence of such delays, the model can generate lead-lag relationships across units because of the leaky dynamics, which induce temporal filtering and make the neuronal activity depend on the past history of the inputs. Incorporating explicit transmission delays would introduce an additional mechanism for generating lead-lag relationships, likely giving rise to richer asymmetric structure in the cross-covariance. Understanding the interplay between these two mechanisms represents an interesting direction for future work. In particular, such an approach could enable a more detailed characterization of the V1–V2 cross-covariances (Fig. 7d–f), in which positive and negative asymmetries emerge at different lags and may reflect qualitatively distinct types of interactions. Importantly, incorporating transmission delays would benefit from also adopting a model capable of faithfully representing fast spiking dynamics beyond the simplified rate model considered here. Our choice of a rate model is motivated by experiments showing that across-area cross-covariances predominantly express slower timescales than within-area ones [62, 6, 8], likely reflecting increased reverberation of activity through indirect pathways. Fast interactions arising from spiking dynamics and direct synaptic transmission have been the focus of complementary theoretical work, studying activity cross-covariances in single areas [63, 64]. Second, our models are characterized by linear dynamics, which greatly simplifies the analysis [64, 65]. Despite their simplicity, linear models can generate rich computational dynamics [66, 34, 67, 68] and can provide accurate approximations of neural dynamics at both local [69] and brain-wide scales [70]. Consistent with this literature, we find that non-trivial directionality properties can emerge even in these simplified networks. Cortical dynamics, however, can deviate from linearity. In particular, nonlinear mechanisms such as gain modulation can change the effective connectivity across different operating regimes, potentially altering the resulting directionality. A possible route toward incorporating these effects in our framework is to consider multiple operating points and local linearizations across epochs [71, 72]. Third, variability in our models arises from fluctuations in external input sources; extending the framework to internally generated variability – such as that produced by spiking dynamics [73, 74, 75] – will be an important direction for future work. Finally, the circuit architecture considered here, although deeply motivated by biology, represents only one of several possible ways of modeling multi-area cortical networks. In our models, cortical areas are composed of locally balanced excitatory and inhibitory populations densely connected through excitatory long-range projections [32]. Alternative models could rely instead on sparse or modular connectivity [76], or architectures in which within- and across-area connections differ in rank [77]. Whether the mechanisms identified here extend to these classes of networks remains an open question for future work.

Our re-analysis of V1–V2 experimental data highlights that, by moving beyond the dominant latent signals and cross-covariances [13], and considering also computationally relevant modes that explain smaller fractions of variance, it is possible to derive simple models that are qualitatively consistent with both the data and the known anatomical organization. Further work will be required to bring these simplified models closer to biological realism and to test their predictions more extensively on experimental data. Of particular interest will be determining whether the distinct timescales expressed in cross-covariance asymmetries can be linked to different modes of signal transmission (e.g., direct versus indirect synaptic pathways), and whether changes in operating point induced by gain modulation between stimulus-evoked and spontaneous activity is sufficient, alone, for inducing shifts in directionality. Finally, extending our analysis to recordings from awake animals, which may exhibit different cross-covariance structures, will be important for assessing the broader relevance of the model.

Our work contributes to the long-standing effort to relate structure and function in neural circuits, and is conceptually aligned with a broad literature from human neuroscience that examines this relationship at the brain-wide scale [78, 79]. Some of these studies have employed functional connectivity measures that rely on temporal asymmetries in the activity statistics [24, 80, 81, 23], and are therefore closely related to the metrics employed here. In this context, our analysis offers two main insights. First, under the assumption of local balance, functional connectivity is expected to relate more closely to the anatomical organization of excitatory than inhibitory subcircuits. Second, assuming that low-resolution imaging signals predominantly reflect global, feature-unspecific latents, the directionality of these signals is related to the overall strength of global signaling targeting a given area. To address brain-wide networks, extending our framework to circuits comprising more than two areas is a crucial next step. This extension is necessary to disentangle the contributions of direct and indirect connectivity loops, which are ubiquitous in cortical architectures, and to explicitly model the origin of input signals carrying internal and cognitive variables, here treated as external. Moving beyond pairs of areas will also clarify how the hierarchy of latent variables identified in this work – framed here as feature-unspecific versus feature-specific components – generalizes to more complex circuits, such as those with higher-dimensional and area-specific tuning properties. Together, these developments provide a principled path toward generating quantitative and mechanistic predictions at the scale of brain-wide networks.

## Methods Circuit model

Throughout the manuscript, we consider recurrent neural networks of *N* units, with activity denoted by ***r***∈ ℝ ^*N*^ . This obeys continuous-time, linear stochastic dynamics defined as:

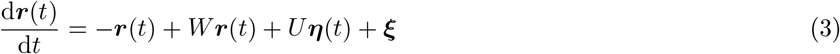

where *W* and *U* ∈ ℝ^*N ×N*^ are matrices that represent, respectively, the recurrent and external input connectivity. The components of the external input vector ***η***(*t*) are modeled as independent stochastic Gaussian processes. For simplicity, we assume that these processes have vanishing mean (stationary nonzero means would affect the activity mean but not its covariance, Equation 2, leaving all results unchanged). For the same reason, we set all components of the bias ***ξ*** to zero. We further assume that ⟨***η***(*t*)***η***(*s*)^*⊤*^ ⟩ = *δ*(*t* −*s*)*I*, where angular brackets denote averages across input realizations, and *I* is the identity matrix. The covariance of the external input is thus given by:

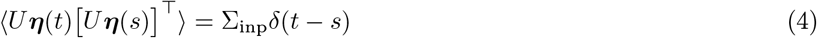

where we defined Σ_inp_ = *UU*^*⊤*^.

By denoting each column vector of *U* as ***u***^*s*^, and each component of ***η*** as *η*_*s*_, Equation 3 can be rewritten as Equation 2. We refer to each independent process *η*_*s*_ as an input source. In general, *N* different input sources can exist. However, we mainly consider cases where most columns of the input matrix *U* vanish, meaning that only a subset of input sources is fed into the circuit at a given time. In these cases, both *U* and Σ_inp_ are rank-deficient. We further assume that input signals are dynamic: within different epochs of a single trial, the identity of the active input sources can change. Mathematically, this is implemented by varying which columns of *U* are nonzero in each epoch.

Equation 3 defines a multivariate Ornstein-Uhlenbeck process [82, 83, 84]. We focus our analysis on stationary activity, that is formally reached in the limit *t*→ ∞, provided that all the eigenvalues of *W* have real part smaller than 1. In the stationary regime, cross-covariance functions (Equation 1) do not depend on time.

### General theory of input-dependent directionality in recurrent circuits

In this section, we establish the theoretical framework used to predict the temporal properties of cross-covariances (Equation 1) generated by recurrent network models. We start by deriving general analytical expressions for cross-covariance functions, and establish conditions under which they exhibit temporal symmetry or asymmetry.

We start defining the propagator *G*(*t*) = exp[(*W* − *I*)*t*], which obeys:

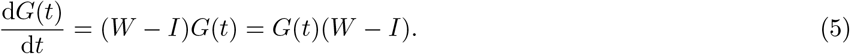

Using the propagator, the solution to Equation 3 can be written as:

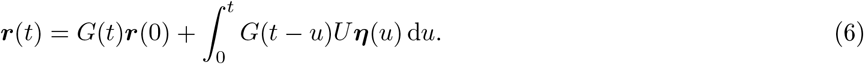

For each input realization, each entry of ***r***(*t*) is normally distributed. The mean of these distributions is given by:

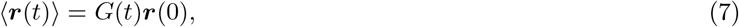

which vanishes in the stationary state. The time-resolved covariance reads instead

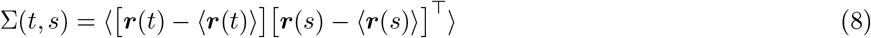

where we assume *t* ≤ *s*, with *s* = *t* + *τ*, and *τ* ≥ 0. From Eqs. 6 and 7, we obtain

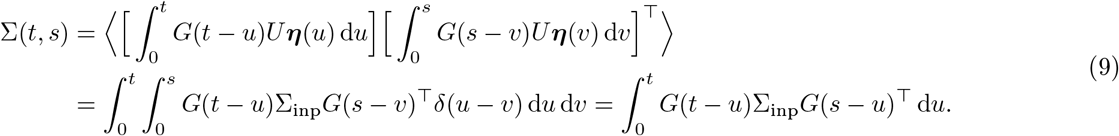

### Properties of the equal-time covariance

The equal-time covariance reads

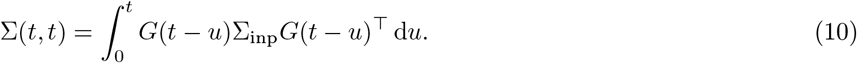

By computing the derivative of this expression with respect to time we find

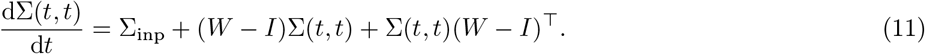

The stationary equal-time covariance, hereafter denoted Σ_0_, is therefore determined by:

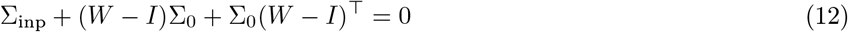

which is typically referred to as the Lyapunov equation. A closed-form analytical solution for this equation exists only in a few special cases (see below).

### Properties of the time-resolved covariance

Consider now the covariance Σ(*t, s*). By taking the derivative of Equation 8 with respect to *τ*, it is possible to verify that Σ(*t, s*) obeys:

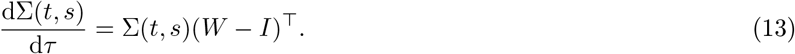

Therefore, integrating over time, we can write:

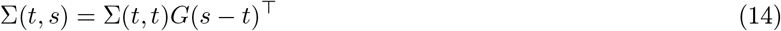

which is also called *regression theorem* in the physics literature [83].

In the stationary state, we can rewrite this expression as:

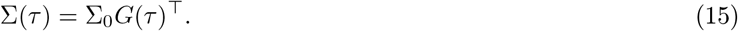

Importantly, because of stationariety, we need to have:

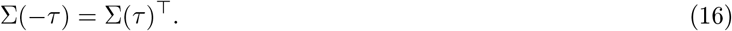

This follows from the requirement that swapping the order of neurons *i* and *j* when computing their covariance should result in flipping the covariance function along the time lag axis: Σ_*ij*_(*t, t* + *τ* ) = Σ_*ji*_(*t, t* − *τ* ).

### Temporal symmetry of covariances

The covariance function is temporally symmetric if:

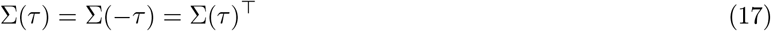

for every time lag *τ* [84, 36, 23]. This condition is also referred to as *equilibrium*, or *time reversibility*, in the physics literature. Combining Eqs. 16 and 15, we see that this is equivalent to imposing

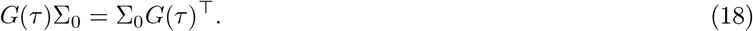

Using the power series representation of *G*, and only considering *τ* small, this condition becomes [83]

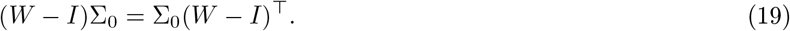

It is straightforward to see that enforcing Equation 19 guarantees that Equation 18 is satisfied at every order in *τ*, because Equation 19 implies

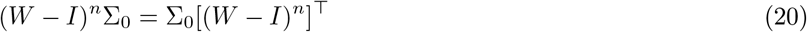

for all values of *n*. To see why this holds for *n* = 2, notice that

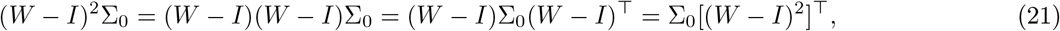

where, in the second and third steps, we applied Equation 19 twice. The relationship can be verified for higher orders of *n* in a similar fashion.

We can now make use of the Lyapunov equation (Equation 12) to further simplify the symmetry condition (Equation 19). Applying Equation 19 onto Equation 12, and solving for Σ_0_, we obtain:

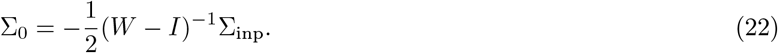

Inserting this into Equation 19, we can rewrite our symmetry condition as

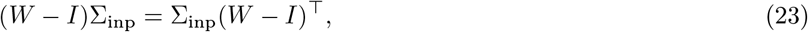

or also

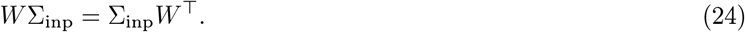

This formulation is particularly useful because it expresses a condition for temporal symmetry solely based on model parameters (the synaptic connectivity and the external inputs covariance). When the stochastic inputs are unstructured and uncorrelated, so that Σ_inp_ = *I*, this condition implies that the synaptic connectivity matrix *W* is symmetric. [36] showed that another class of matrices that satisfies Equation 23 is given by *W* = Σ_inp_*D*, with *D* diagonal. This connectivity matrix depends on the statistics of the external inputs via a Hebbian-like structure.

### Changes of basis

Suppose now we analyze the network dynamics in a different reference system, defined by a linear transformation of coordinates. The relationship between activity in the original and transformed systems is given by

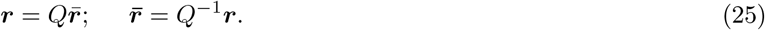

The dynamics in the new reference system read:

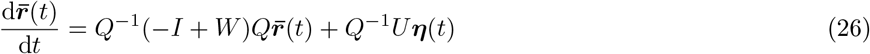

which can be re-cast in the usual form:

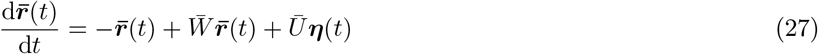

by defining the transformed connectivity matrix 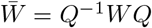. = *Q*^*−*1^*WQ* and inputs *Ū* = *Q*^*−*1^*U*, so that 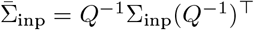 [84]

The temporal symmetry condition in the new reference system reads:

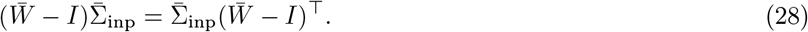

By using the expressions for the transformed connectivity and inputs covariance, it is easy to show that the equation above is equivalent to Equation 23. This makes sense, as global aspects of the temporal dynamics should not depend on the reference system choice. This also implies that we can study the temporal symmetry properties of activity in a given neural circuit by using equivalently the neural space and the transformed space.

### Eigenvector basis

We now focus on a specific basis, given by the eigenvectors of the connectivity matrix *W* . As we shall detail in the rest of the section, this basis comes with two key advantages: (i) it allows us to derive a more intuitive interpretation of the temporal symmetry condition (Equation 23); (ii) it allows us to derive closed-form analytical solutions for the covariance functions.

We therefore assume that the connectivity matrix is diagonalizable, with eigenvectors {***p***_*k*_} _*k*_ and associated eigenvalues {*λ*_*k*_} _*k*_. We denote with *P* the matrix constructed by stacking horizontally the column vectors corresponding to the eigenvectors. (For real symmetric *W*, the eigenvectors are real and orthonormal, so the eigenvector matrix *P* satisfies *P*^*−*1^ = *P*^*⊤*^. For diagonalizable, non-normal *W*, the columns of *P* and the rows of *P*^*−*1^—the right and left eigenvectors, respectively—generally differ; *P* maps eigenmode coordinates to neural coordinates, while *P*^*−*1^ performs the reverse transformation. The right eigenvectors specify spatial activity patterns, while the overlap of the left eigenvectors with the input determines how strongly each mode is driven). In the eigenvector basis (obtained from Equation 25 by setting *Q* = *P* ), the transformed connectivity matrix 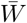 is diagonal, namely: 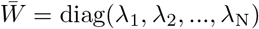.

The temporal symmetry condition (Equation 24) reads:

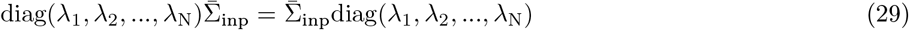

i.e. the transformed connectivity and covariance input matrices commute. We now assume that all eigenvalues are distinct: *λ*_*i*_ *≠λ*_*j*_ ∀*i, j* (but see next section for discussion of a more general case). Under this condition, Equation 29 holds if and only if the transformed input covariance is diagonal, i.e.

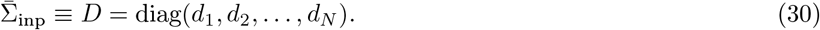

Equivalently, temporal asymmetry holds if the transformed inputs in the eigenvector space have zero cross-covariance (but can have different variance). In the following section, we provide an alternative and more intuitive derivation of this condition.

Equation 30 directly follows from the following lemma: let *A* and *D* be two *N N* matrices, with *D* diagonal and distinct entries. Assume that those matrices commute: *AD* = *DA*; we can then prove that *A* is also diagonal. We have:

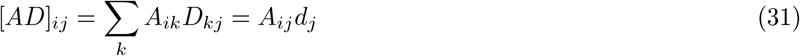

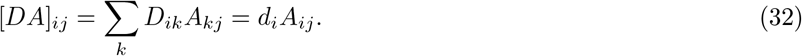

Therefore:

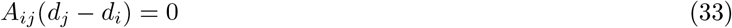

implying *A*_*ij*_ = 0 for *i* ≠ *j*.

### Computing activity covariance in the eigenvector space

We can make use of the eigenvector basis to derive explicit analytical expressions for the covariance function Σ(*t, s*). This covariance can be computed from the transformed covariance as

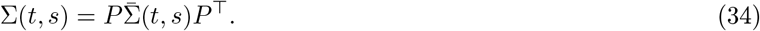

We can therefore focus on computing 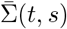, which can be done by evaluating Equation 9 in the eigenvectors space. In the stationary regime, we have

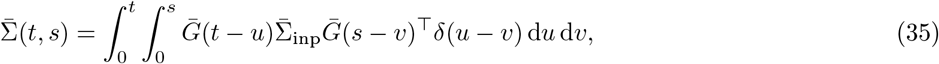

where we have defined the transformed propagator

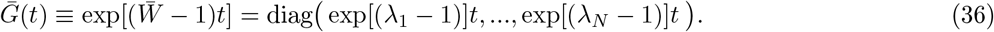

We once again assume *t* ≤ *s*. We can integrate over the Dirac delta function, to obtain

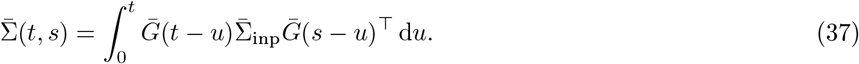

We now use that, for every matrix *A*, we have: [*D*^1^*AD*^2^]_*ij*_ = [*D*^1^]_*ii*_[*A*]_*ij*_[*D*^2^]_*jj*_ when *D*^1^ and *D*^2^ are diagonal matrices. We get:

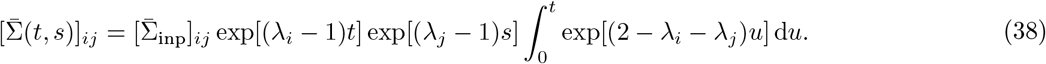

Solving the integrals, we obtain:

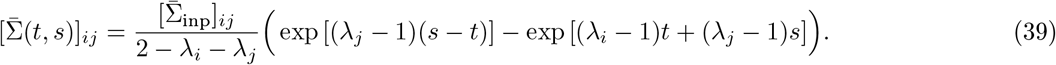

Making use of the stationariety assumption (*t, s* → ∞), this simplifies to:

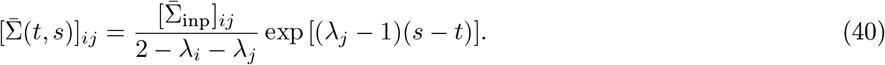

One specific implication of this expression is that the variance of activity along each eigenvector direction is given by:

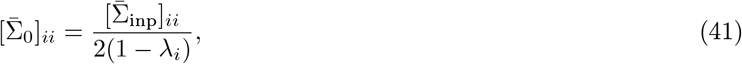

implying that variance is only amplified along eigenvector directions associated with positive eigenvalues.

In the stationary regime, we can further rewrite Equation 40 as a function of the delay *τ* = *s* − *t* ≥ 0. We have:

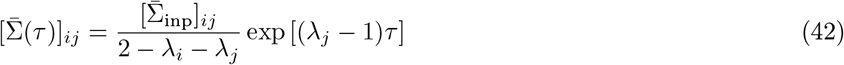

while, by using Equation 16

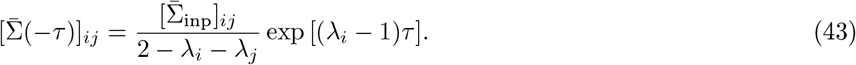

Note that the covariance function in the eigenvector space takes a simple shape: an exponential decay with different characteristic timescales for positive and negative *τ* (Fig. 3d). Throughout this work, we use these expressions, together with Equation 34, to evaluate theoretical estimates for cross-covariances (Fig. 3, Fig. 4, Fig. 6, Supplementary Fig. 1–8).

Under which condition is this covariance matrix temporally symmetric? Imposing 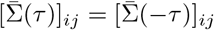 yields

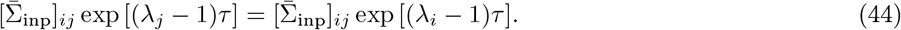

We again assume that eigenvalues are distinct. For the diagonal entries, *i* = *j*, Equation 44 is always verified. For the off-diagonal entries, *i* = *j*, Equation 44 is only verified when the off-diagonal entries of the transformed input covariance vanish: 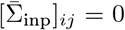. We therefore retrieve the symmetry condition derived earlier (Equation 30): the transformed covariance 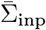 must be diagonal.

We can now provide an intuitive explanation for the condition we just derived. The eigenvector basis is particularly suitable for studying the temporal properties of activity because, in this basis, each activity component is characterized by a unique timescale, set by the eigenvalue. In particular, since eigenvalues differ, each activity component corresponds to an input signal filtered at a different timescale. If the input signals associated with different eigenvectors are correlated, the emergent activity along these directions will also be correlated, but will exhibit distinct timescales. Cross-correlating these activities (either in the eigenvector, or in the neural space) gives rise to lead-lag behavior and therefore temporally asymmetric covariance functions. If the input signals associated with different eigenvectors are uncorrelated, the emergent activity along these directions will also be uncorrelated. Cross-correlating these activities does not generate any lead-lag behavior, and results in temporally symmetric covariance functions. (Those vanish in the eigenvector space, but not in the neural space, see Fig. 3).

In Fig. 3, we illustrate this intuitive reasoning in a simplified scenario where inputs are one-dimensional. In that case, input signals associated with different eigenvectors are uncorrelated if and only if the input vector is aligned with one of the eigenvectors.

Finally, note that if a pair of repeated eigenvalues exists, the corresponding entries of the transformed covariance are temporally symmetric. However, the remaining entries remain asymmetric. Consequently, temporal asymmetries are still expected among many, if not all, pairs of units in the original neural space. More generally, as long as the connectivity matrix has at least two distinct eigenvalues, temporal asymmetry can be expected in the neural space. One specific example is given by rank-one connectivity matrices for which one eigenvalue is non-zero, and all the remaining eigenvalues vanish. As detailed later on, these matrices can give rise to temporally asymmetric cross-covariances.

### Conditions for temporal symmetry in eigenvector space

Under which conditions on the recurrent connectivity *W* and the input matrix *U* the transformed inputs in the eigenvector space are uncorrelated? To answer this question, we need to determine when

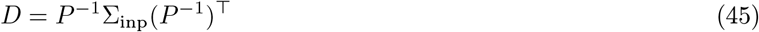

or, equivalently,

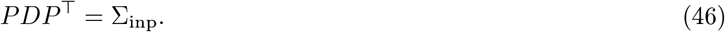

By noticing that Σ_inp_ = *UU*^*⊤*^, we can rewrite this as

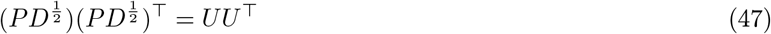

which has the solution

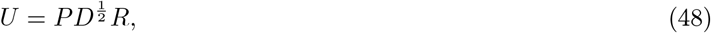

where *R* is any orthogonal matrix obeying *R*^*⊤*^*R* = *I*.

For fixed connectivity matrix and fixed eigenvectors *P*, Equation 48 determines a constraint on the input matrix *U* .

This equation has a straightforward interpretation: the input transformation *U* that guarantees temporal symmetry is obtained by scaling the eigenvectors of the connectivity matrix through arbitrary values (given by the diagonal entries 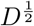), followed by an additional distance-preserving, orthogonal transformation (given by *R*, applied row-wise). Since *D* and *R* are not fixed, for a given *P*, a family of solutions exists for *U* . Example inputs from this family of solutions are illustrated in Supplementary Fig. 2.

Alternatively, Equation 48 can equivalently be rewritten as:

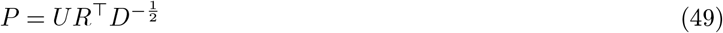

which expresses a constraint on the connectivity eigenvectors *P* for a fixed input matrix *U* . (Note that we assumed here that all diagonal entries of *D* are non-vanishing).

To understand the origin of the degeneracy in Eqs. 49 and 48, we can resort to a geometrical intuition. We know that the distribution of stochastic inputs in the neural space follows a multivariate normal distribution, with covariance matrix Σ_inp_. This distribution arises from a linear projection, defined by *U*, of a spherical multivariate normal distribution with covariance matrix *I* (for the variable ***η***, Equation 3).

When determining the set of connectivity eigenvectors that ensure time symmetry (Equation 49), we seek a linear transformation *P*^*−*1^ that maps the multivariate normal distribution with covariance Σ_inp_ back into a uncorrelated one. (In fact, Equation 45 can be interpreted a whitening transformation [85]). One possible solution to this problem is choosing 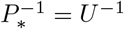, which maps the inputs back to the original variable ***η*** – and, therefore, to a spherical distribution. After projecting to the original space, we can then stretch each dimension independently, yielding the uncorrelated family of solutions:

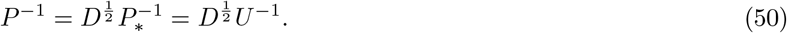

Crucially, due to symmetry of the original spherical distribution, any solution that applies an orthogonal transformation before stretching is also valid. This leads to the general solution:

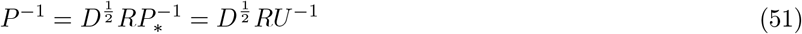

from which Equation 49 can be retrieved.

One could observe that another solution 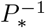 can be naturally constructed from the principal components of the multivariate normal distribution. In that case, we would start with the eigenvector decomposition of Σ_inp_:

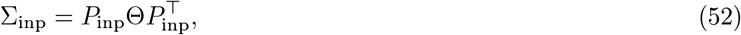

where Θ is diagonal and *P*_inp_ is orthogonal. We would then set: 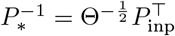. This solution is in fact just one of infinitely many others that can be obtained by starting from 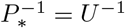 and applying an orthogonal transformation *R*, as discussed above. In this case, we have 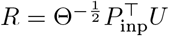, which can be showed to be orthogonal.

### Low-dimensional inputs for temporal symmetry

We note that, according to Equation 48, the input matrix *U* can be singular. Specifically, its rank satisfies:

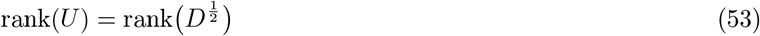

implying that *U* has a rank smaller than *N* when some diagonal entries of *D* vanish. In such cases, the input matrix is determined by only a subset of the eigenvectors, and the external inputs are low-dimensional.

We start examining a solution in which all but the first diagonal element of *D* vanish. In this case, Equation 48 yields:

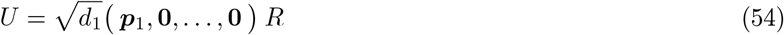

where ***p***_1_ is the first eigenvector of *W* and **0** represents a column vector of zeros. This expression can be rewritten as:

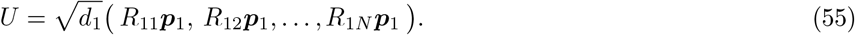

The total input to the network is one-dimensional, and can be re-expressed as ***p***_1_*η*_eff_(*t*), where *η*_eff_ is just a rescaled noise source:

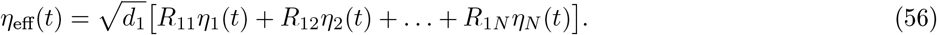

We conclude that one-dimensional inputs satisfying Equation 48 always involve an input vector that is aligned with one of the eigenvectors of *W* . This is because the terms *d*_1_ and *R* in Equation 54 do not qualitatively affect the input configuration, but just rescale the input variance.

Next, we consider a solution in which all but the first two diagonal elements of *D* vanish. In this case, we have:

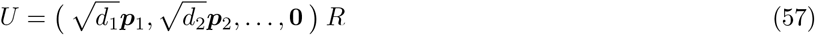

which expands to:

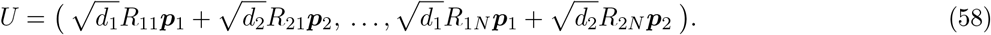

The total input can now be expressed as ***p***_1_*η*_eff, 1_(*t*) + ***p***_2_*η*_eff, 2_(*t*), where

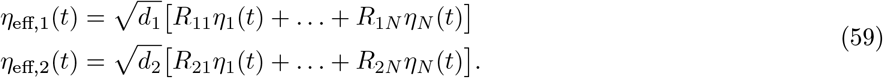

These two effective input sources are generally correlated (unless *R* = *I*, see paragraph below), with their correlation structure controlled by *d*_1_, *d*_2_, and *R*. Varying those degrees of freedom, we therefore obtain in this case a family of qualitatively different input configurations.

### Eigenvector-aligned solution

We discuss a particular solution to Equation 48, obtained by setting *R* = *I*, which gives

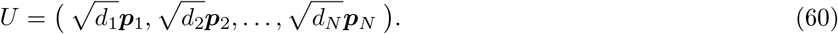

In this case, each input source (or, equivalently, each column of *U* ) is associated with a single eigenvector. This solution has a special property: one can independently modulate the diagonal entries of *D*, parametrizing the strength of each input source, without affecting the directional properties of the cross-covariances. In particular, one can tune a given input source to be either off (by setting the diagonal entry to 0) or on (by setting the diagonal entry to a non-zero value), implementing a circuit configuration where different input sources contribute to activity at different time epochs. In all scenarios, the cross-covariances remain temporally symmetric.

By controlling the number of the diagonal entries of *D* that are non-zero, one can build solutions in the form of Equation 60 of any dimensionality. For example, a one-dimensional input can be constructed as:

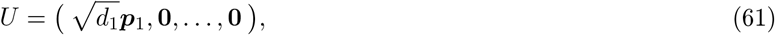

for which the total input is equal to 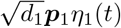. Note that this solution is qualitatively identical to Equation 55. One example circuit endowed with this type of input is illustrated in Fig. 3, violet lines. A two-dimensional input can be instead constructed as:

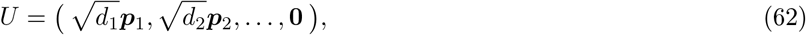

for which the total input is equal to 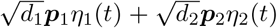. This solution represents a specific case within the broader two-dimensional family described by Equation 58. One example circuit endowed with this type of input matrix is illustrated in Supplementary Fig. 2.

### Activity-input covariance

So far, we have focused on activity covariances. However, it is also informative to compute covariances between neural activity and inputs, as this provides insight into how input signals are transformed by the network (see Supplementary Fig. 3B). Specifically, we consider the covariance:

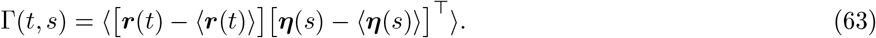

Using Equation 6, in the stationary regime we have:

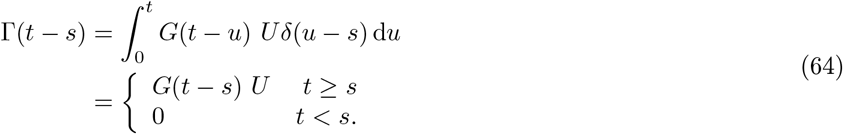

To evaluate this expression, we use again the eigenvector basis. We start from 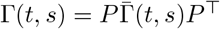 and then compute 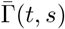. Using Equation 36, we have

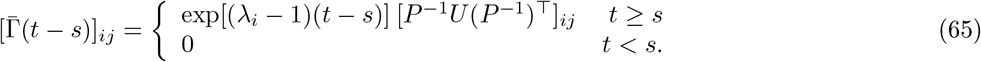

### Rank-one connectivity

We study in detail a specific class of synaptic connectivity matrices that can be written as a rank-one matrix [86, 68]:

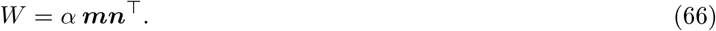

Here, *α* is a scalar and ***m*** and ***n*** are unit-norm vectors. The only non-zero eigenvalue of *W* is given by *λ* = *α* ***m***^*⊤*^***n***, and is associated with the eigenvector ***m***. The remaining eigenvectors are all orthogonal to ***n***. Our interest in this type of matrix arises from the fact that some simple excitatory-inhibitory circuits can be recast in this form. We begin by discussing the results obtained so far in the context of rank-one matrices. We then show how, for this specific connectivity choice, the same results can be derived in a more direct manner, leading to equations that are easier to interpret.

The condition for temporal symmetry (Equation 24) reads in this case:

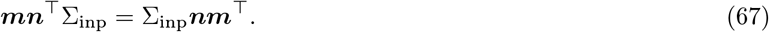

We further know that temporal symmetry is ensured for inputs satisfying Equation 48. Here, the columns of the eigenvector matrix *P* are given by the vector ***m***, and *N*− 1 linearly independent vectors chosen to be orthogonal to ***n***. This highlights that, in the context of a rank-one connectivity matrix, most input directions preserve temporal symmetry. (Intuitively, this is because most input directions do not engage recurrent dynamics and therefore cannot trigger input filtering across multiple timescales). The only input direction that promotes temporal asymmetry is one that has a nonzero component along ***n***, while not being fully aligned with ***m***.

For rank-one matrices, we can also derive a closed-form solution for the covariance function. We start from the propagator, which in rank-one matrices takes a simple form:

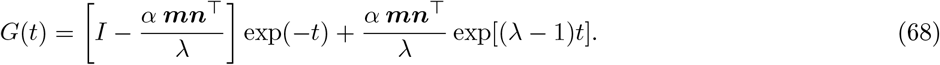

(Note that this specific expression is only well-posed for *λ ≠*0). Plugging this expression into Equation 6, and focusing on the stationary state, we have

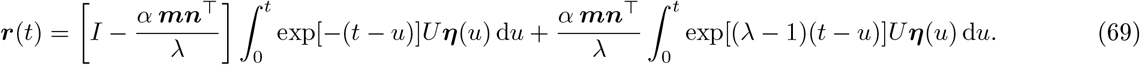

This equation shows that activity consists of two terms, each with dynamics arising from the filtering of input noise at different timescales.

We set *τ* = *s* − *t >* 0. We can use Equation 69 to directly compute the covariance function as

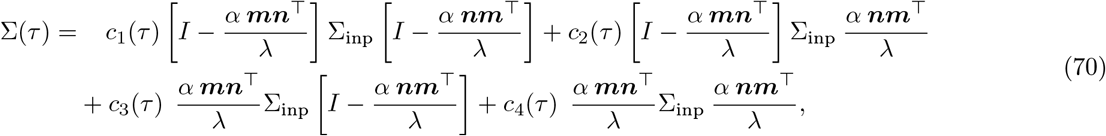

where the time-dependent coefficients can be computed assuming stationariety. We have:

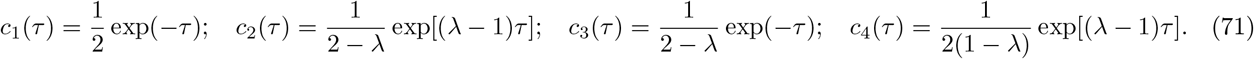

Temporal symmetry requires Σ(*τ* ) = Σ(− *τ* ), with Σ(−*τ* ) = Σ(*τ* )^*⊤*^. From Equation 70, it is evident that only the second and third terms of the covariance are non-symmetric and, therefore, can introduce temporal asymmetry. From the intuitive understanding developed thus far, this makes sense, as these are the terms that cross-correlate the parts of activity associated with different timescales (Equation 69). Specifically, we can rewrite Equation 70 as:

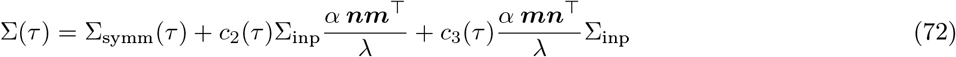

where Σ_symm_(*τ* ) = Σ_symm_(*τ* )^*⊤*^. The condition for temporal symmetry therefore becomes:

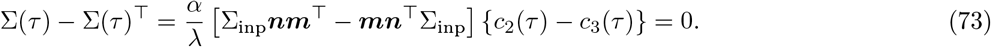

Since *c*_2_(*τ* ) ≠ *c*_3_(*τ* ) when *λ* = 0 (distinct eigenvalues), we have that the term in square brackets has to vanish, and therefore Equation 67 is retrieved.

The fact that the cross-covariance function can be computed in closed form for this specific type of synaptic connectivity allows us to make an important observation: as the connectivity strength *α* increases, the amplitude of the temporally symmetric component of the covariance grows faster than the asymmetric one. This is because the symmetric part scales linearly with *α*, while the asymmetric part does not. To see this, recall that *λ* = *α* ***m***^*⊤*^***n***, and note that we can rewrite

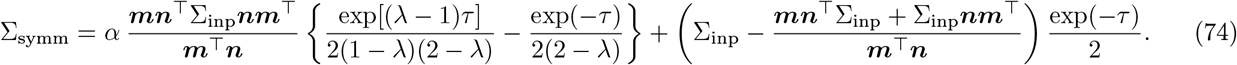

Note also that, when ***m***^*⊤*^***n*** *>* 0, increasing *α* brings the network closer to instability: *λ* approaches 1, at which point *c*_4_(*τ* ) diverges, while *c*_2_(*τ* ) and *c*_3_(*τ* ) remain finite. Taken together, these observations indicate that strong temporal asymmetry – unlike input amplification effects, see [87] – arise in regimes with intermediate recurrent connectivity, far from instability.

### Single-area excitatory-inhibitory model

Up to this point, our derivations have focused on generic circuits (Equation 3) with arbitrary synaptic connectivity *W* . From here onward, we turn to connectivity matrices of specific biological relevance and examine how external inputs shape their cross-covariance structure. As a starting point, we consider a model of a single cortical area composed of one excitatory (E) and one inhibitory (I) population [39, 34]. This analysis serves as a building block for the analysis of the two-area model discussed in the main text.

In this model, the activity vector (Equation 3) can be expressed as

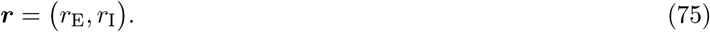

The synaptic connectivity matrix is given by:

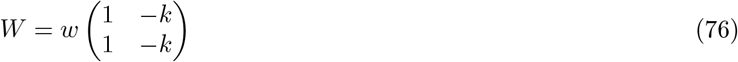

where *w* and *k* are two positive scalars encoding, respectively, the total synaptic strength and relative inhibition dominance. The eigenvectors of *W* are given by:

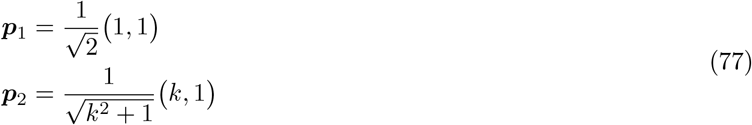

corresponding, respectively, to eigenvalues *λ*_1_ = *w*(1− *k*) and *λ*_2_ = 0. We focus on inhibition-stabilized networks, for which *w*≥ 1 and *k*≥ 1. Exact excitation-inhibition balance in the synaptic connectivity is reached for *k* = 1.

This matrix is rank-one, and can be rewritten as in Equation 66 by setting

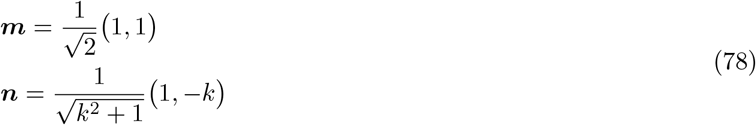

as well as 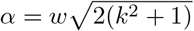.

As long as the connectivity is approximately balanced (i.e., *k* is not much larger than 1), the eigenvector ***p***_2_ substantially overlaps with ***p***_1_, and both are nearly orthogonal to the vector ***n***. The two eigenvectors encode a direction along which excitatory and inhibitory populations co-fluctuate, whereas ***n*** encodes a direction along which these populations are modulated in opposition [34].

### Cross-covariance asymmetry

Under which input configurations do we expect directional versus non-directional interactions between the E and I populations? We begin by noting that, since the connectivity matrix is non-symmetric – as is the case for all biologically-inspired matrices considered from now on – high-dimensional, unstructured inputs (Σ_inp_ = *I*) are generally expected to induce directional interactions. We then narrow our focus to one-dimensional inputs in the form

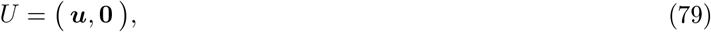

where ***u*** = *u*_E_, *u*_I_, and the total input to the network is given by ***u****η*_1_(*t*). We assume that this input originates from another brain area and enters the modeled circuit via excitatory synapses. Accordingly, we restrict our analysis to the case where *u*_E_, *u*_I_≥ 0.

Our analysis so far indicates that symmetric cross-covariances arise when the input vector ***u*** is aligned with one of the eigenvectors of the connectivity matrix. According to Equation 77, this occurs when ***u*** is approximately aligned with the balanced direction, i.e., when excitatory and inhibitory units receive similar input (Supplementary Fig. 3 and 4). In this regime, E and I populations act as filters characterized by a single timescale, and recurrent dynamics do not enrich the temporal structure of the input. As a result, fluctuations in E and I activity are temporally synchronized. In contrast, strong directional interactions arise when ***u*** is poorly aligned with the balanced direction. Previous work has shown that such inputs (which differentially target E and I neurons) give rise to substantial non-normal input amplification [34, 87, 68]. Therefore, the same input directions that drive non-normal amplification also generate pronounced temporal asymmetries in the cross-covariance.

As discussed in a earlier section, we expect temporal asymmetry in the cross-covariance to be maximal at intermediate coupling strengths, *w* (see also Supplementary Fig. 4a). This corresponds to a regime in which synaptic current balance is not enforced to be tight, and inhibition lags behind excitation, resulting in a sluggish tracking [88]. Previous studies have shown that the amount of non-normal amplification increases instead with the coupling strength [87, 68]. Therefore, while temporal asymmetry and non-normal amplification are both maximized for the same input direction, they peak at different coupling values.

### Time window of integration

The direction of the input vector ***u*** influences not only cross-covariance asymmetry and input amplification, but also the effective dynamics of the network response – specifically, how quickly the E and I populations react to inputs and the timescales at which they filter them. When ***u*** aligns with the balanced direction, it is approximately orthogonal to the connectivity vector ***n***. In this configuration, recurrent connectivity is minimally engaged [86], and inputs are primarily filtered by the intrinsic neuronal leak, resulting in a relatively fast response. This is reflected in the activity–input cross-covariance (Supplementary Fig. 3b beige), which decays rapidly at negative lags. The response can become even faster when ***u*** is slightly biased toward stronger I than E input, though this effect reverses if the imbalance becomes too large (see below). In contrast, when ***u*** aligns with the non-balanced direction, recurrent connectivity plays a central role in shaping the response, via the non-normal amplification mechanism [34]. This amplification comes at the cost of a slightly slower response, as evident in the more gradual decay of the activity–input cross-covariance (Supplementary Fig. 3b red and blue). Importantly, differences in these cross-covariances are confined to short negative lags and decay at longer lags. This indicates that the differences in integration dynamics are transient, consistent with the fact that the eigenvalues of the recurrent connectivity (and, therefore, the asymptotic timescales of the dynamics) remain unchanged across input directions.

### The over-inhibited regime

According to our analysis so far, strong temporal asymmetry and input amplification can co-occur when inputs are aligned with non-balanced directions. Those directions can express inputs that predominantly target either the excitatory (*u*_E_≫ *u*_I_ ≥0) or the inhibitory (*u*_I_ ≫*u*_E_ ≥0) population. While both types of inputs promote temporal asymmetry and non-normal amplification [34], they correspond to qualitatively different dynamical regimes. We describe these two regimes below.

When inputs predominantly target the excitatory population (Supplementary Fig. 3a, red), both temporal asymmetry and amplification are strong. Both E and I units are positively correlated with the external input, as showed by activity–input covariances. The E unit leads the I unit in time (the delay can be interpreted as the time required for I to generate a negative copy of the input received by E) [89]. Because both populations track the same input source, their cross-covariance is positive at all lags.

When inputs predominantly target the inhibitory population (Supplementary Fig. 3a, blue), temporal asymmetry remains strong, but non-normal amplification is weaker. In this regime, the external input generates a transiently large activity in the I unit, which exerts a strong suppressive effect on E activity. The E unit becomes anti-correlated with the input, as it can only sustain high activity when the net inhibitory influence from the input via the I population is minimal. This behavior leads to negative cross-covariances between E and I activities at negative lags. The activity-input covariance for the I unit displays a positive peak at small negative delays due to the direct influence of the external input, followed by a negative tail at larger negative delays, which reflects the delayed, negative copy of the input received by the E unit (a phenomenon related to the so-called *paradoxical effect*, [39]). Because the E–I cross-covariance in this regime includes both positive and negative lobes, with their relative amplitudes displaying high sensitivity to parameter values, measures of asymmetry (in particular, the asymmetry score) display large values and high variability. We name this regime the over-inhibited regime.

Networks operating in the over-inhibited regime typically satisfy *u*_I_ ≥*u*_E_ ≥0 (Supplementary Fig. 4). The exact bounds, however, depend on the network’s connectivity parameters. Practically, to determine whether a model lies in the over-inhibited regime, we compute the activity–input covariance of the E unit (Equation 63) and check whether it becomes negative for some negative time lag.

In this work, we focus on networks operating outside the over-inhibited regime. This choice is motivated by several considerations. First, both theoretical [89] and experimental [90, 91] studies suggest that cortical networks operate in a regime where excitatory and inhibitory neurons co-fluctuate in response to external drives – a mechanism that supports asynchronous spiking even under shared input. Second, although strong feedforward inhibition is observed in cortical circuits, there is, to our knowledge, no experimental evidence that it induces dynamics resembling those of the over-inhibited regime. It has been shown that feedforward inhibition from thalamus to sensory cortex sharpens the temporal responses of excitatory neurons [40, 29, 41]. This aligns with the behavior of our model outside the over-inhibited regime: as *u*_I_ increases from 0 to values approaching *u*_E_, the activity–input cross-covariance decays more rapidly. By contrast, the over-inhibited regime would involve even stronger inhibition, leading to responses that are not only brief but slow and suppressive. Third, in scenarios where inputs are the main driver of cortical activity (as is often the case in sensory areas), the strong inhibition typical of the over-inhibited regime would likely lead excitatory neurons below threshold. In such conditions, predictions from our linear model would lose validity, and cross-covariances would appear flat and temporally symmetric.

Finally, we remark that the strength of synapses mediating incoming inputs to excitatory and inhibitory populations can be measured experimentally [40, 29, 92]. These studies have reported strong synapses for both excitatory and inhibitory inputs, with relative strengths varying with stimulus properties [29], as well as cortical areas and layers [93]. Work by [92, 93] suggests that, within the cortical hierarchy, feedforward interactions involve relatively stronger inhibitory inputs than feedback interactions, leading to faster dynamics. Thus, just as there is no clear experimental indication of the over-inhibited regime in feedforward thalamocortical interactions (see paragraph above), there is likewise no strong evidence for such a regime in cortical feedback pathways.

### Two-areas excitatory-inhibitory model

We now consider a four-dimensional model as in Equation 3 consisting of two coupled E-I populations, where each population corresponds to a distinct cortical area and inter-areal connections are exclusively excitatory (Fig. 4a). Ordering units by area, we define the activity and input vectors as

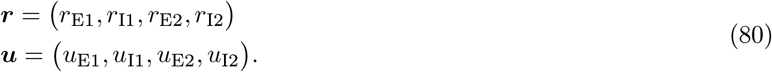

We focus our attention on a specific synaptic connectivity matrix:

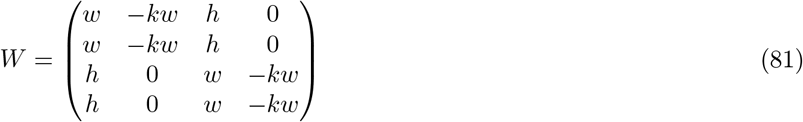

where *h* is a positive scalar controlling the strength of across-area connections. We also use the notation:

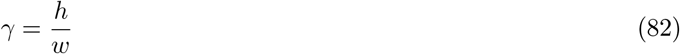

to indicate the relative strength of across-area connections with respect to within-area ones.

This matrix embodies several simplifying assumptions: (i) within-area connectivity is identical across the two areas; (ii) inter-areal connectivity is symmetric in the feedforward (first →second area) and feedback (second→ first) directions; (iii) inter-areal projections target both E and I populations in the recipient area with equal strength. We use this construction to avoid making assumptions about the detailed biological structure of mesoscopic cortical connectivity (which are often poorly constrained by available data, and strongly differ across theoretical models [66, 94]). In addition, this construction ensures stability across a broad parameter range, and affords analytical tractability due to its low rank (rank-two; see below). To evaluate the robustness of our results, we also analyze alternative connectivity matrices in which one or more of these simplifying assumptions are relaxed (see Supplementary Figs. 6 and 7).

### Connectivity eigenvectors

This synaptic connectivity admits two eigenvectors given by:

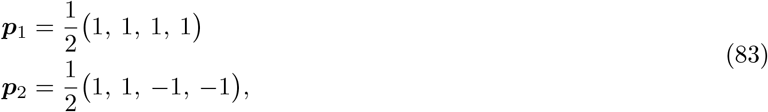

associated with eigenvalues

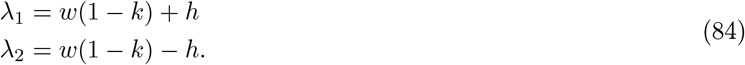

For almost balanced within-area connectivity (*k*≃ 1), these two eigenvalues are equal to± *h*. Across-area connectivity therefore gives rise to a positive and a negative eigenvalue (and, correspondingly, to a slow and a fast activity timescale).

The positive eigenvalue reflects positive feedback across areas, mediated by excitatory inter-areal projections, and corresponds to an activity pattern where the two areas co-fluctuate. The negative eigenvalue is associated instead with broken positive feedback, and an activity pattern where the two areas fluctuate in opposition. The fact that *λ*_1_ *>* 0 *> λ*_2_ suggests that a large fraction of the activity variance in this circuit is expected along ***p***_1_ (see Equation 41), implying that cross-covariances among all pairs of units are in general positive-valued [32].

Note that across-area connectivity generates an instability, approximately, at *h* = 1. Motivated by the observation that cortical areas can operate in a fast dynamical regime, here we focus on values of *h* that are far from the instability. We also assume *h < w*.

The remaining two eigenvectors are associated with eigenvalues *λ*_3_ = *λ*_4_ = 0. Those eigenvectors are unspecified, and form a two-dimensional family. Two specific eigenvectors within this family are given by

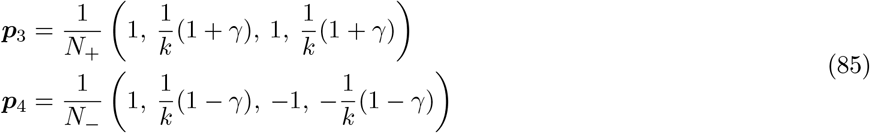

where

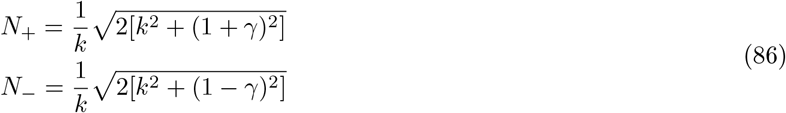

are two normalization constants. Starting from those eigenvectors, we can parametrize the entire eigenvectors family as

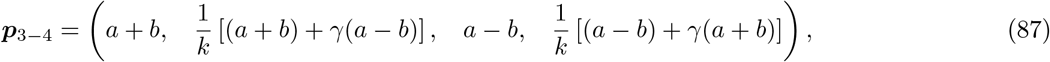

with *a* and *b* real-valued.

Our results so far indicate that when the input vector of a one-dimensional input source is aligned with one of the eigenvectors, the resulting cross-covariances among all circuit’s units are temporally symmetric. For the first two eigenvectors, ***p***_1_ and ***p***_2_, this can be understood intuitively. In these cases, the inputs to E and I units within each area are aligned along the balanced direction, leading to synchronous activity between local E and I populations. The E and I populations across the two areas are further synchronous due to the model built-in symmetries (namely, identical connectivity and input strengths across the two areas). The eigenvectors ***p***_3_, ***p***_4_, as well as the broader family defined by Equation 87, are perhaps more interesting. They correspond to input vectors for which fully symmetric cross-covariances can be obtained even if the inputs to local E and I populations are not identical, and the input strengths are not symmetric across areas. These eigenvectors are analyzed more in detail in the section below.

We conclude this section by providing explicit expressions for the matrix *P*, which collects all eigenvectors as columns, and for its inverse. These matrices will be used in the eigenvector analysis presented later. We have:

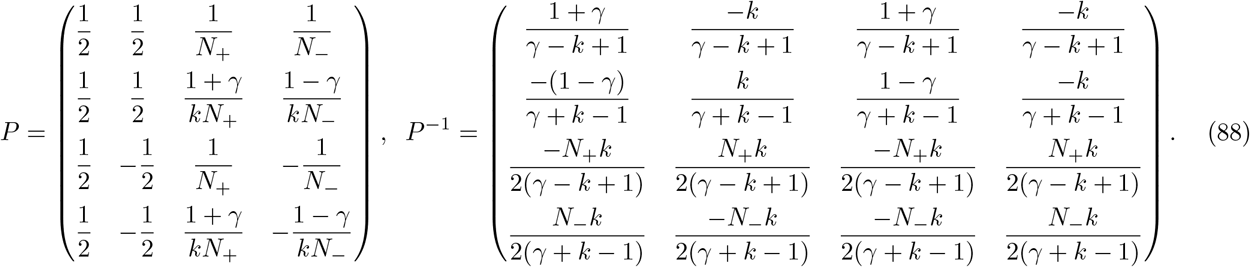

**Analysis of specific eigenvectors**

Before presenting an in-depth analysis of the excitatory–excitatory cross-covariance, we briefly outline a few properties of the eigenvector family defined in Equation 87 which facilitate the interpretation of the results presented in Fig. 4. This eigenvector family encompasses vectors where the excitatory (or, alternatively, inhibitory) components are identical across both areas. To identify the eigenvector with identical excitatory inputs across areas, we set the first and third components of Equation 87 to an arbitrary value *u*_E_, yielding:

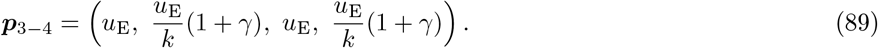

This eigenvector is symmetric across areas but exhibits asymmetry between local E and I units. By aligning the input vector with this eigenvector, one can therefore achieve temporally symmetric cross-covariances between local E and I populations even if inputs to E and I are unequal. The value of the input to I that ensures temporal symmetry depends on the connectivity parameters, and is maximized for large *γ* and for *k* = 1. In general, for almost balanced within-area connectivity, this value exceeds *u*_E_.

We have discussed that, in single-area models, inputs to I that are stronger than those to E drive the circuit into an over-inhibited regime, characterized by highly directional E–I interactions (see also Supplementary Fig. 3, 4). Equation 89 indicates that local E–I interactions can remain non-directional even in the presence of higher I inputs. This occurs because across-area connectivity counteracts over-inhibition by fostering large-amplitude activity across both areas via excitatory feedback. This effect is maximized for balanced local connectivity (*k* = 1), which enhances input amplification (Supplementary Fig. 4b). Input configurations with identical excitatory inputs are discussed in Fig. 5b. In those plots, the value of the input to I predicted by the second and fourth entries of Equation 89 is depicted as a black dot. That value signals, approximately, the maximum input to I for which, thanks to across-area connectivity, activity remains outside the over-inhibited regime (dark area).

To find the eigenvector with identical inhibitory input cross areas, we set the second and fourth components of Equation 87 to an arbitrary value *u*_I_, and obtain:

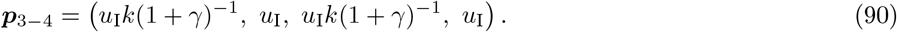

In this case, the value of the input to E that guarantees temporal symmetry is minimized for large *γ* and for *k* = 1. Input configurations with identical inhibitory inputs are discussed in Fig. 5a. In those plots, the value of the input to E predicted by the second and fourth entries of Equation 90 is depicted as a black dot and signals, approximately, the minimum input to E for which activity remains outside the over-inhibited regime.

### Eigenvector-based analysis of excitatory-excitatory cross-covariance

In our analysis so far, we have used the eigenvectors of the synaptic connectivity matrix to identify input directions that give rise to synchronous activity and temporally symmetric cross-covariances across all units in the circuit. We now narrow our focus to just two units – specifically, the excitatory populations in each area – and examine how their cross-covariance and the directionality of their interactions depend on the input vector. This focus is motivated by the fact that excitatory cells constitute the majority of cortical neurons and are more frequently the target of experimental recordings. We note that, due to the fully symmetric connectivity between the two areas, and the sign reversal symmetry of the network dynamics (Equation 3), any input vector with entries: ***u*** = (*u*_E_, *± u*_I_, *u*_E_, *± u*_I_ )leads to a temporally symmetric excitatory-excitatory cross-covariance. Our goal is therefore to characterize directionality for input vectors that do not display such a symmetry.

We start expressing activity in the two E populations from their eigenvector components. Using Eqs. 83 and 85, we can write

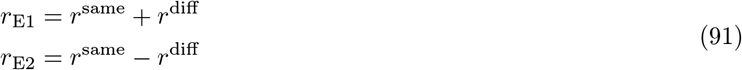

where the signals *r*^same^ and *r*^diff^ are defined as:

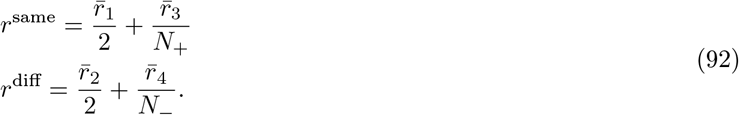

These two signals coincide with the eigenvector signals illustrated in Fig. 5a–b. Both *r*^same^ and *r*^diff^ are constructed from two out of the four components that represent activity in the eigenvector space. The signal *r*^same^ includes the slowest component 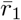, associated with the eigenvalue *λ*_1_, as well as the neutral component 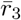, corresponding to a zero eigenvalue. In contrast, *r*^diff^ contains the fastest component 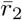, associated with the eigenvalue *λ*_2_, and the neutral component 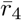. Equation 91 indicates that co-fluctuations in the activity of the two areas are captured by the signal *r*^same^, which expresses the slowest timescales. In contrast, the signal with fast timescales, *r*^diff^, captures the component of activity that varies in opposition across the two areas [32].

Using Equation 91, the excitatory-excitatory cross-covariance can be expressed as

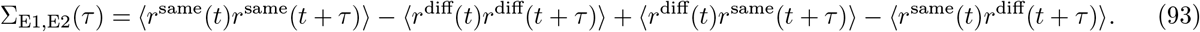

The first two terms represent auto-covariances and are thus symmetric with respect to the time lag *τ* . In contrast, the last two terms correspond to cross-covariances, which may be asymmetric; their difference yields an antisymmetric function in *τ* . This antisymmetric component underlies the temporal asymmetry of the cross-covariance and therefore governs the directionality between the two excitatory populations. Crucially, if this antisymmetric term is small relative to the symmetric one, the cross-covariance will be approximately symmetric. This can occur, in principle, when the amplitude of *r*^same^ is small relative to *r*^diff^, or vice versa. In the parameter region we focus on, because of the positive feedback mediated by excitatory inter-area connectivity, *r*^same^ has larger amplitude than *r*^diff^. Therefore, Σ_E1,E2_ is in general a positive function, which is approximately symmetric if the amplitude of *r*^diff^ is small with respect to *r*^same^. In order to compute *r*^same^ and *r*^diff^, we need to compute the components of activity in the eigenvector space, 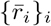.

Using Equation 26 with *Q* = *P*, we see that these evolve according to

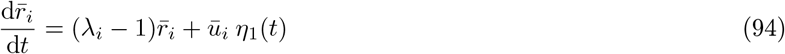

where we defined ***ū*** = *P*^*−*1^***u***. We introduce the shorthand notation

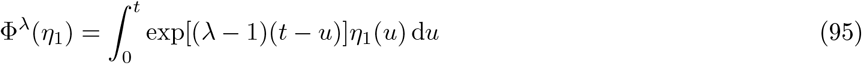

for the filtered input *η*_1_ at timescale *λ* 1. We then obtain, in the stationary regime: 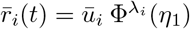. From these expressions, the signals *r*^same^ and *r*^diff^ (Equation 92) can be computed as:

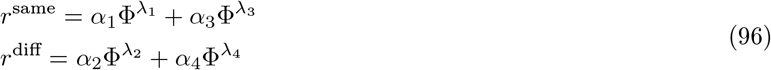

where the coefficients {*α*_*i*_}_*i*_ are given by

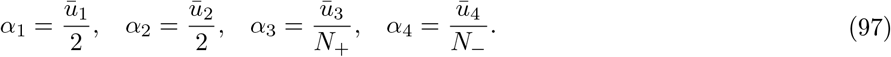

To compare the relative amplitudes of *r*^same^ and *r*^diff^, we need to evaluate the coefficients *α*_*i i*_, which depend on the choice of the input vector.

#### Inputs to a single area

We begin by analyzing a simple setup, where the one-dimensional input is provided exclusively to the first of the two areas. This corresponds to an input vector of the form:

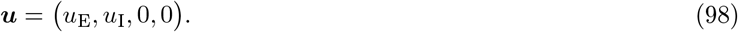

Using Equation 97 together with Equation 88, we obtain:

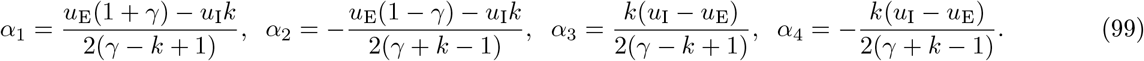

We analyze these coefficients in two simplified cases. First, we consider the case where the external input equally feeds into the excitatory and inhibitory units (*u*_E_ = *u*_I_ = 1). The coefficients simplify to:

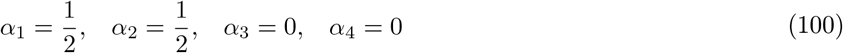

so that, from Equation 96:

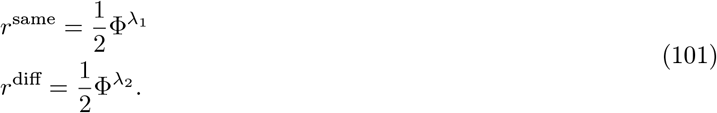

The amplitude of the two filtered signals 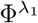 and 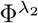 depends on the eigenvalues *λ*_1_ = *w*(1 −*k*)+*h* and *λ*_2_ = *w*(1 −*k*) − *h*. In the regime we are interested in, for which within-area connectivity is almost balanced (*k*≃ 1), and the across-area connectivity is not too strong, both *λ*_1_ and *λ*_2_ are not too far from zero, and the two filtered signals have comparable magnitude. Therefore, *r*^same^ and *r*^diff^ are expected to have similar amplitude, implying asymmetric cross-covariances and directional interactions.

In agreement with the results discussed in Fig. 4, we can further verify that directionality is oriented from the first area to the second one. This is because the cross-covariance can be re-expressed as

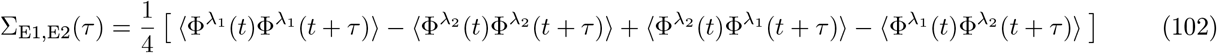

where terms in the right hand side obey:

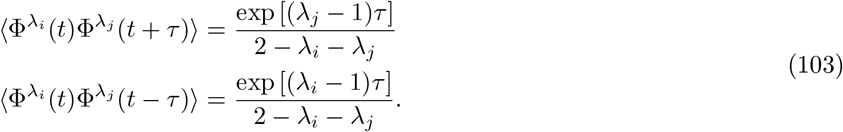

By noting that *λ*_1_ *> λ*_2_, one can easily verify that the sum of the first two terms in Equation 102 give rise to a positive symmetric function, while the sum of the last two to an antisymmetric function with positive asymmetry.

We can further consider another specific configuration, in which the input targets only the excitatory unit (*u*_E_ = 1, *u*_I_ = 0). We get:

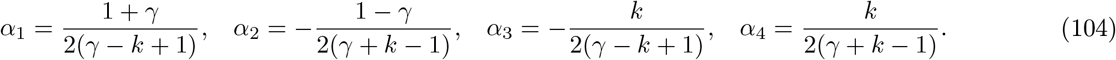

As in previous case, the coefficients associated with *r*^diff^, *α*_2_ and *α*_4_, do not vanish. However, the signal *r*^diff^ could still be suppressed with respect to *r*^same^ if *α*_2_ and *α*_4_ had similar magnitudes and opposite signs (note that this would also require the two filtered signals 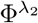 and 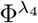 to have similar amplitude and temporal structure, see next paragraphs). This would however entail:

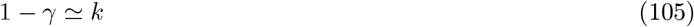

which is never satisfied, as the left- and right-hand sides are, respectively, always smaller and larger than one. In the limit case where the equality holds (*k* = 1 and *γ* = 0), also *α*_1_ and *α*_3_ have similar amplitude and different sign; therefore, also *r*^same^ has small amplitude, and temporally symmetric cross-covariances cannot be expected. All in all, also for this input configuration, asymmetric cross-covariances and directional interactions are expected.

#### Identical inputs to I

We now consider input configurations where both areas receive non-zero input. We start considering the setup illustrated in Fig. 5a, where the input provided to the I populations is identical. This corresponds to an input vector of the form:

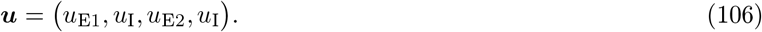

The coefficients in this case are:

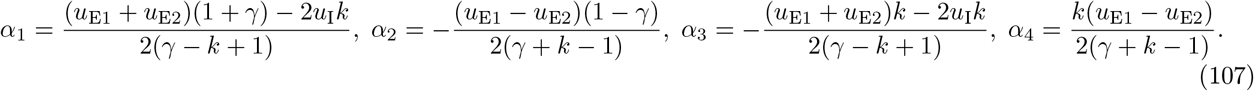

We know that when *u*_E1_ = *u*_E2_, the model is fully symmetric across areas, and the cross-covariance must be symmetric. Indeed, in that case, we find that *α*_2_ = *α*_4_ = 0, implying that *r*^diff^ vanishes, and activity in the two excitatory populations is fully synchronous.

In the general case, *r*^diff^ could still have a small amplitude, provided that *α*_2_ and *α*_4_ have similar magnitude but opposite signs. However, this condition leads back to Equation 105, which is never satisfied. Therefore, for this type of inputs, cross-covariances are generally expected to be temporally asymmetric.

It is instructive to estimate Equation 107 in the limit of large *γ*, corresponding to a regime where the strength of inter-area connectivity, *h*, is much larger with respect to within-area one, *w*. This limit deviates significantly from the parameter choices employed in this work (for which *w > h*), but is useful to clarify the role of across-area connectivity in shaping directionality. In this limit, the inhibitory populations are effectively disconnected from the circuits, and the circuit in which the two excitatory populations are embedded is effectively two-dimensional. Keeping only the dominant order in *γ* in Equation 107, we get

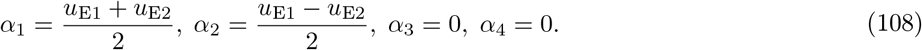

This case is similar to the one analyzed in Equation 100. Following the same analysis steps, if we assume that the strongest input is fed into the first area, *u*_E1_ *> u*_E2_, we can conclude that the cross-covariance is temporally asymmetric, with the first area leading the second one.

#### Identical inputs to E

We then consider the setup illustrated in Fig. 5b, where the input provided to the E populations, but not the I populations, is identical across areas. This corresponds to an input vector of the form:

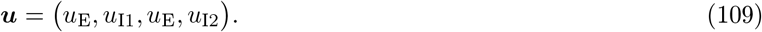

The coefficients are given by:

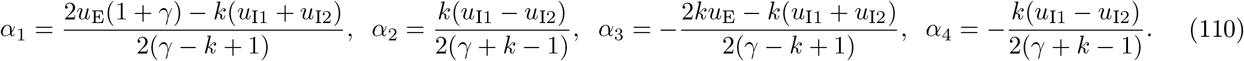

Similarly to the previous paragraph, when *u*_I1_ = *u*_I2_, the signal *r*^diff^ vanishes, and the cross-covariance is symmetric. For generic input values, the coefficients in Equation 110 behave differently than those in Equation 107. In particular, for all parameter values, *α*_2_ and *α*_4_ are characterized by equal magnitude but opposite sign, leading to an approximate cancellation in the second row of Equation 96, and thus to a potentially small *r*^diff^ amplitude. This equality in magnitude does not hold for *α*_1_ and *α*_3_, implying that the amplitude of *r*^same^ cannot be expected to be small.

Under which conditions is the amplitude of *r*^diff^ effectively suppressed? We can rewrite

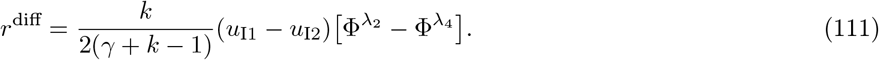

In order for this signal to be effectively small, both the difference term 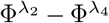 and the prefactor term *k/*(*γ* + *k*− 1) must remain small. We start analyzing the difference term 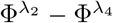. Since *λ*_2_ = *w*(1 −*k*) −*h* and *λ*_4_ = 0, for fixed *h* and *w*, the difference is minimized when *k* = 1. Furthermore, for *k* = 1, the difference is minimized when the across-area connectivity parameter *h* is small. Note that, when this is verified, *r*^diff^ is not only characterized by small amplitude, but also by strongly reduced temporal structure (see, for instance, the bottom panel of Fig. 5b, where the temporal structure is much weaker compared to Fig. 5a).

We now consider the prefactor term *k/*(*γ* + *k* −1). This term displays a fairly shallow dependence on *k*, supporting *k* = 1 as a suitable choice for promoting cancellation. (As discussed earlier, this choice also reduces the extent of the over-inhibited regime, further favoring small temporal asymmetries.) For *k* = 1, the ratio *k/*(*γ* + *k*− 1) in Equation 111 however diverges as *γ* approaches zero. This indicates that too small across-area connectivity is ineffective at suppressing the *r*^diff^ signal.

Combining the two pieces of the analysis together, we conclude that optimal cancellation occurs in a regime where *h* is small but nonzero, and *w* is not too strong with respect to it, so that *γ* = *h/w* takes finite values. In other words, the approximate cancellation of *r*^diff^ is optimal when inter-area connectivity is weak, yet still not negligible with respect to within-area connectivity. These dependencies are verified in Supplementary Fig. 6 and 7.

Finally, we consider again the limit of large *γ*. For the input configuration discussed in this paragraph, the limit yields a two-dimensional circuit consisting of excitatory units that receive identical input. By symmetry, a symmetric cross-covariance is expected. Indeed, we have

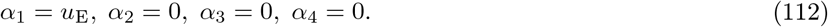

The signal *r*^diff^ vanishes (as could also be expected from the fact that the prefactor in Equation 111 converges to zero). As a final remark, the analysis so far focused on the activity of the two excitatory populations and their cross-covariance. However, it can be readily extended to include cross-covariances between excitatory-inhibitory and inhibitory-inhibitory population pairs. Such an extension reveals that the cancellation observed for inputs of the form given in Equation 109 is specific to the excitatory-excitatory cross-covariance. Also, it can be easily showed that this cancellation critically depends on the excitatory nature of the inter-area connectivity, as it does not occur in circuit models that also incorporate inhibitory inter-area projections.

### Small-*h* analysis

After using the eigenvector perspective to build intuition about cross-covariances and their directionality, we now discuss a complementary mathematical approach. This approach assumes that the across-area connectivity, *h*, is weak, and analyzes the resulting activity using a power expansion that retains only the dominant terms. In particular, we set

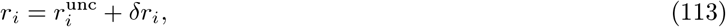

where the first term on the right-hand side corresponds to activity in the absence of across-area connections (*h* = 0, uncoupled term), and the second one (*h >* 0, coupled term) is linear in *h*. Higher-order terms in *h* are neglected.

Setting *h* = 0, we see that the uncoupled terms obey:

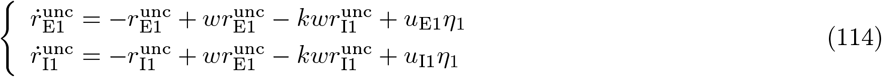

and similarly for the second area. Uncoupled activity thus results from the processing of external inputs by single-area E-I circuitry. Our results so far indicate that the amplitude and temporal structure of this activity are governed by the strength of input to E and I populations (see also Supplementary Fig. 3 and 4). In particular, the amplitude of activity increases with input strength (Supplementary Fig. 4c).

The coupled activity term obeys instead:

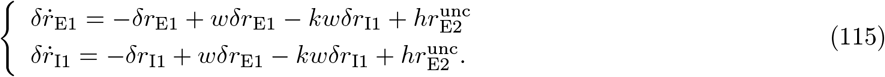

We can again think of this system of equations as describing the local activity of area 1 in response to external inputs – however, in this case, the inputs 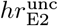 are provided by the other area. These inputs are not temporally white, as their dynamics result from the temporal integration of the external noise (Equation 114).

We use Eqs. 114 and 115 to make two important observations. First, the amplitude of coupled activity in one area scales with the amplitude of the uncoupled activity in the other. In particular, if uncoupled activity is smaller in the first area than in the second, then the coupled activity in the first area is larger than in the second. As a net effect, the relative contribution to activity due to across-area coupling is larger in the first area. This phenomenon is rooted in the excitatory nature of connectivity between areas, which acts to redistribute variance across the circuit. Second, the coupled activity terms, representing the contributions to activity that stem from across-area connections, are generally slower than the uncoupled terms. This is a consequence of the fact that the temporal filtering expressed by Equation 115 can only slow down signals.

The latter observation is, strictly speaking, only valid under the assumption of weak inter-areal connectivity. However, we expect it to hold (at least qualitatively, and for most circuit configurations) also in the regime of intermediate coupling. This is because, as *h* increases, the dominant eigenvalue of the synaptic connectivity matrix, *λ*_1_, becomes larger. The across-area dynamics represented by coupled activity terms therefore express increasingly slow timescales. This implies that the timescale separation between the uncoupled and coupled activity components observed in the small *h* regime becomes even more pronounced at stronger inter-area coupling.

In the following, we examine the implications of the small-*h* analysis for the three input scenarios discussed in the previous section.

#### Inputs to a single area

We start again discussing the case of inputs fed into a single area (Equation 98). Since 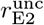 vanishes, we have:

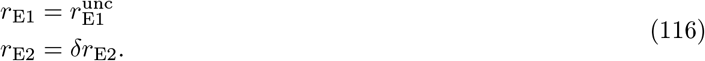

Thus, the first area follows approximately uncoupled dynamics, while the second is dominated by across-area signals. In particular, activity in the second area is given by *δr*_E2_, which is a temporally filtered version of activity in the first one, 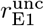 (Equation 115). Therefore, the second area lags behind the first. Note that interactions are directed from the area of high variance to area of low variance, and are aligned with the effective flow of inputs.

#### Identical inputs to I

We consider now the case of inputs characterized by identical strength to I populations (Equation 106). For simplicity, we focus on one specific input configuration, for which the dynamics falls outside of the over-inhibited regime:

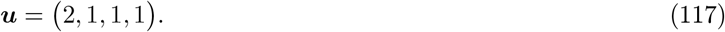

We start by analyzing the uncoupled activity (Equation 114). The inputs to the first area are such that the local dynamics amplify the input, leading to large activity and relatively slow input responses (see Supplementary Fig. 3, 4). In contrast, the inputs to the second area do not undergo amplification, resulting in smaller activity and faster responses. Therefore, in the absence of across-area coupling, the first area would lag behind the second (see Fig. 4h). We then consider the coupled activity (Equation 115). This contribution is much stronger in the second area, which becomes dominated by slow inter-area signaling and starts lagging behind the first one (Fig. 5a). For sufficiently large values of *h*, this contribution dominates the cross-covariance. As in the previous case, directionality is therefore directed from the area of high variance, to the area of low variance.

#### Identical inputs to E

We consider now the case of inputs characterized by identical strength to the E populations (Equation 109). We consider the configuration:

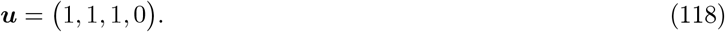

As for Equation 117, the difference in input across areas for this configuration is equal to 1.

The uncoupled activity for this input behaves opposite of the previous configuration: activity in the first area is small and fast, while activity in the second area is large and slow. As a consequence, in the absence of across-area coupling, the first area precede the second one (Fig. 4h). Coupled activity, carrying slow dynamics, is stronger in the first than in the second area. Also in this case, therefore, across-area connectivity tends to overcome the local directional properties of the circuit. In contrast to the previous case, however, this compensation mechanism is weaker: instead of reversing directionality, it leads to non-directional interactions. To understand why this is the case, we can consider the variance unbalance driving the reversal in the two scenarios.

Across the two scenarios, the uncoupled variance in the low-activity areas is identical. However, the uncoupled variance in the high-activity areas is not, as it emerges from inputs (2,1) and (1,0). In particular, the (2,1) configuration generates larger variance than (1,0). This is because, whenever the input to the E population is increased by a given amount, in order to maintain the same output activity, the input to I has to be increased by a larger amount (Supplementary Fig. 4c grey dots indicate the input values considered here). In other terms, the activity variance in E-I circuits is more sensitive to increases in the input to E than to I populations.

We remark that, in both scenarios, directionality is tightly linked to the properties of non-normal amplification performed by excitatory-inhibitory circuits [34]. For uncoupled areas, non-normal amplification determines directionality by setting different internal timescales. For coupled areas, it specifies the variance unbalance, and therefore the effective flux of the slow signal across the two areas.

### Modified two-areas excitatory-inhibitory model

We now consider variations of the model described at the beginning of this section. In particular, we relax some of the assumptions embedded in the synaptic connectivity matrix, Equation 81. The resulting models are used in Supplementary Fig. 6 to investigate the robustness of our main findings. In Supplementary Fig. 6c left, we relax the assumption that within-area connectivity is identical across areas. We replace the parameter *w* by two different parameters, *w*_1_ and *w*_2_, modeling the strength of excitatory recurrence in areas 1 and 2. This is motivated by the gradient in excitatory connectivity strength observed along the cortical hierarchy [95]. In Fig. 4f and Supplementary Fig. 6c center, we relax the assumption that inter-area connectivity is symmetric in the two directions. We replace the parameter *h* by two different parameters, *h*_FF_ and *h*_FB_, to model the strength of inter-areal connectivity from the first to the second area and vice-versa. In Supplementary Fig. 6d right, we relax the assumption that inter-areal projections target E and I populations in the recipient area with equal strength. We introduce an additional parameter *α* to model the strength of across-area connectivity to E and I populations as *h* and *αh*. This circuit variant is motivated by experimental evidence showing differences in the strength of across-area inputs to E and I in visual cortical areas [92]. For this circuit variant, the eigenvalues can become complex; we chose parameters to avoid that regime.

### Feature-specific two-areas model

We now consider an eight-dimensional model as in Equation 3 consisting of two coupled cortical areas, each comprising two pairs of excitatory and inhibitory populations. Each populations pair represents a local subcircuit selectively tuned to a specific stimulus feature. For concreteness, we label these features as A and B.

Ordering units by area and by stimulus selectivity, the activity vector is defined as:

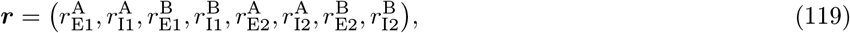

and similarly for the input vector. We define the connectivity matrix *W* as:

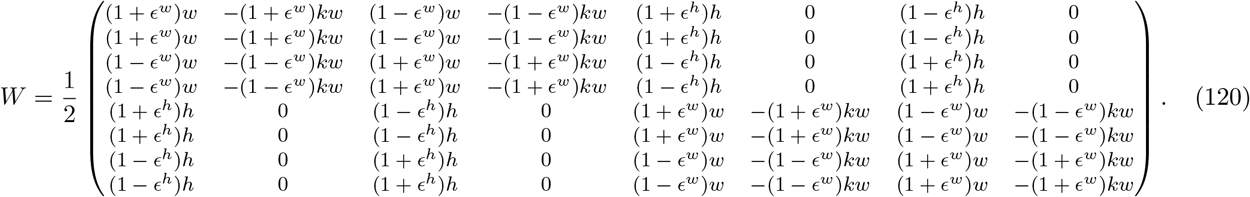

Each feature-specific excitatory-inhibitory subcircuit is locally connected and receives input from the other subcircuit within the same area, as well as from both subcircuits located in the other area. Local connectivity within a subcircuit follows the same structure as in previous models but is scaled by a factor of (1 + *ϵ*^*w*^)*/*2, with 0 *< ϵ*^*w*^ *<* 1. Connections from and to the other subcircuit within the same area have the same form but are scaled by (1 −*ϵ*^*w*^)*/*2. Thus, the parameter *ϵ*^*w*^ controls the degree of feature-specific segregation in within-area connectivity, with *ϵ*^*w*^ = 0 corresponding to no feature specialization, and *ϵ*^*w*^ = 1 corresponding to two completely disconnected subcircuits. Inter-area connectivity is purely excitatory. The strength of these connections is scaled by (1 + *ϵ*^*h*^)*/*2 when connecting subcircuits with the same feature selectivity, and by (1 −*ϵ*^*h*^)*/*2 when connecting subcircuits with different selectivity (0 *< ϵ*^*h*^ *<* 1). Therefore, the parameter *ϵ*^*h*^ controls the degree of feature specialization in inter-area connectivity, with *ϵ*^*h*^ = 0 corresponding to no feature specialization, and *ϵ*^*h*^ = 1 corresponding to fully specialized inter-area connectivity.

We parametrize the input vector as

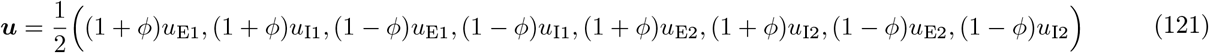

with 0 ≤*ϕ*≤ 1. For *ϕ* = 0, the input vector is fully feature-unspecific: the entries of the E or I units within the same area, with different feature tuning, are identical. In this case, under stationary conditions, the model formally reduces to the simpler two-area system analyzed earlier. This is due to the model’s symmetry, which implies *r*^A^ = *r*^B^ ≡ *r*_*kl*_ for *k*∈ {E, I} and *l* ∈{1, 2 }; the eight-dimensional system thus effectively becomes four-dimensional, with connectivity described exactly by Equation 81.

### Connectivity eigenvectors

This synaptic connectivity admits four eigenvectors given by:

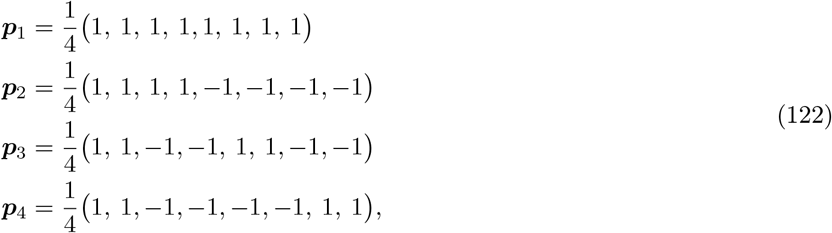

associated with eigenvalues

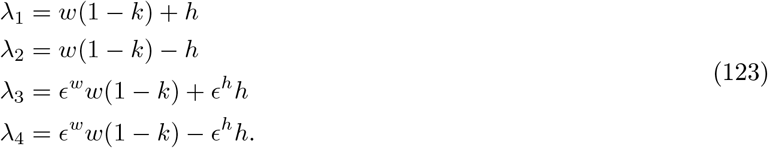

The first two eigenvalues and their corresponding eigenvectors represent feature-unspecific modes, which are either consistent or opposite across areas. For balanced within-area connectivity (*k*≃ 1), the first (resp. second) eigenvector is associated with a positive (resp. negative) eigenvalue. The third and fourth eigenvectors correspond to feature-specific modes, again either consistent or opposite across areas. For balanced within-area connectivity, the third (resp. fourth) eigenvector has a positive (resp. negative) eigenvalue whenever inter-area connectivity is characterized by nonzero specialization (*ϵ*^*h*^ *>* 0).

The remaining four eigenvalues vanish. The corresponding eigenvectors have in general complex expressions. Similarly to the simpler two-areas model, these eigenvectors depend on connectivity parameters, and are non-orthogonal to the first four.

Recurrent processing amplifies activity along eigenvector directions associated with positive eigenvalues (Equation 41). Consequently, in this model, only the eigenvectors corresponding to *λ*_1_ and *λ*_3_ undergo amplification. Because, in general, *λ*_3_ *< λ*_1_, the third eigenvector is amplified less strongly than the first [32], and the two receive equal amplification only in the special case of fully specialized inter-areal connectivity (*ϵ*^*h*^ = 1). These statements are qualitative: in practice, the precise amplification of each eigenvector also depends on the orientation of the input vectors (Equation 41).

### Population structure of activity

Before quantifying across-area directionality, we characterize the population structure of the activity patterns generated by this model. We analyze activity from the perspective of within-area latent signals constructed from excitatory populations. The latents are defined as

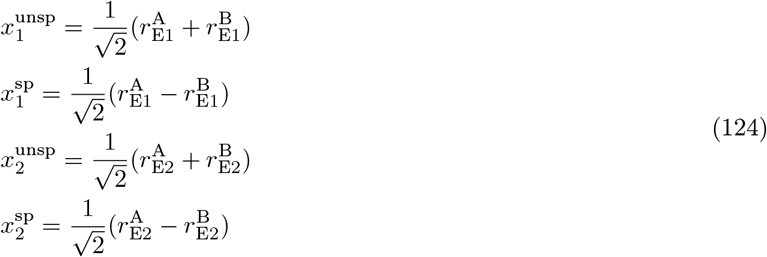

and are obtained by projecting the activity vector ***r*** onto the directions:

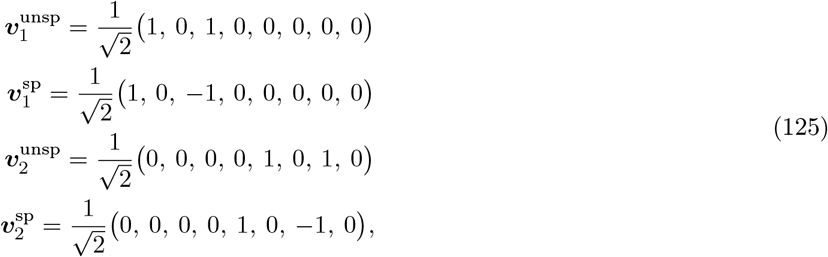

representing orthonormal feature-specific and unspecific directions within the two areas. Note that, in this basis, activating a single excitatory unit within an area (e.g., ***r*** = (1, 0, 0, 0, 0, 0, 0, 0)) leads to activation of both latent signals (e.g., 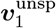 and 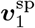). This occurs because, although the activation is purely feature-specific, the non-negativity constraint on activity and the orthogonality of vectors in Equation 125 necessarily introduce a feature-unspecific component.

We define the matrix *V* by stacking the four vectors from Equation 125 as rows. Denoting the latent signals as

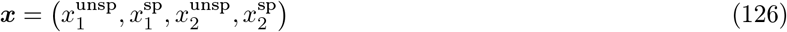

we can express them compactly as ***x*** = *V* ***r***. We next use the eigenvectors of the connectivity matrix, as derived in the previous section, to characterize the temporal structure of the latent signals. We start observing that we can write 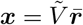, with 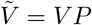. This matrix has the following structure

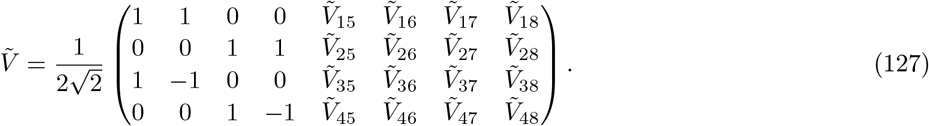

(For conciseness, we did not explicit the values of entries in the last four columns, which are not crucial for this analysis). We conclude that the feature-unspecific latents 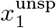 and 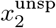 express the eigenvector components 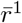 and 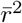, which correspond to the fastest and slowest timescales, respectively. In contrast, the feature-specific latents 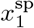 and 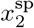 express the eigenvector components 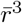 and 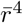, reflecting intermediate timescales. Finally, both specific and unspecific latents express the neutral components from 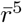 to 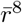.

We have already noted that the two eigenvector components 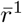 and 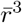 are expected to express the largest variance, due to positive eigenvalues. Equation 127 indicates that among these components, only 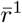 contributes to the feature-unspecific latents, while only 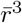 contributes to the specific ones. Because 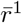 is amplified more than 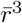, we expect that, in both areas, unspecific latents have larger variance than specific ones [32]. Quantitatively, the magnitude of this difference in amplification depends on model parameters, and is predicted to be largest when across-area connections are fully unspecific (*ϵ*^*h*^ = 0), for which the separation between the eigenvalues *λ*_1_ and *λ*_3_ is maximal.

### Directionality of latent signals cross-covariances

We compute cross-covariances between pairs of feature-specific and unspecific latent signals across areas. As discussed so far, unspecific latents capture the largest share of the variance within each area. Since they are both dominated by the eigenvector component 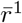, these two latents are also strongly positively correlated. Consequently, we expect unspecific cross-covariances to exhibit the largest amplitude (Fig. 6b–c). To formalize this argument, we evaluate the amplitude of the unspecific and specific cross-covariances at *τ* = 0, restricting the analysis to the activity components 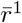 through 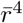, which correspond to non-zero eigenvalues. (Although this is an approximation, it substantially simplifies the derivation without altering the underlying argument.) We obtain:

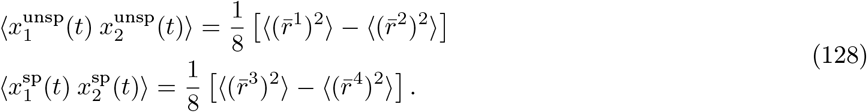

Since 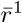 typically has larger amplitude than 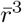, and 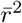 is generally smaller than 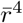, the unspecific cross-covariance is expected to exceed, in amplitude, the specific one. This difference is quantified systematically in Fig. 6d–e. A second expected difference between the unspecific and specific cross-covariances is their temporal decay, which is generally slower in the unspecific case, due to the longer timescale expressed by 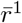 compared to 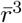 [32].

We now characterize the directionality. Interestingly, this can be predicted by directly leveraging the results previously derived for unspecialized circuits. To see why, we begin by deriving the equations governing the temporal evolution of unspecific latents. Making use of Eqs. 3 and 120, we have that

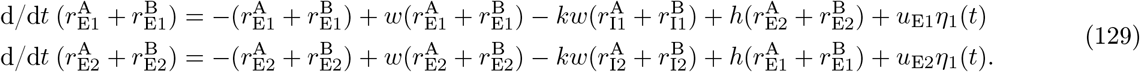

This equation shows that the unspecific activity of excitatory populations (and, by an analogous derivation, inhibitory populations) across the two areas follows the same dynamics as those analyzed for circuits with unspecific connectivity. Building on those results, we conclude that the directionality of unspecific latents is determined by the relative strength of the total inputs to the two excitatory populations, *u*_E1_ and *u*_E2_, while remaining largely insensitive to the total inputs to the inhibitory populations, *u*_I1_ and *u*_I2_. In particular, the area receiving the strongest input leads the other in time. All-in-all, the directionality of the unspecific cross-covariance reflects the total flow of external inputs across areas.

The dynamics of specific latent signals obeys instead

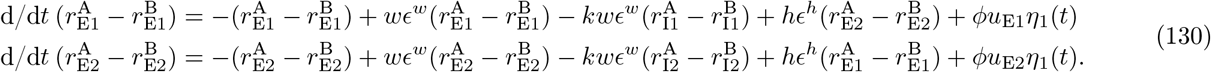

These dynamics are similar to those studied in simpler two-area models with no specialization, but they correspond to a circuit with rescaled connectivity: within-area connectivity is rescaled by *ϵ*^*w*^, and across-area connectivity by *ϵ*^*h*^. When *ϵ*^*w*^ = *ϵ*^*h*^, the recurrent connectivity is proportional to the one analyzed earlier. In that case, specific latents behave qualitatively like unspecific ones: directionality is determined by the relative strength of *u*_E1_ and *u*_E2_, flowing from the more strongly driven area to the other. When *ϵ*^*h*^≫ *ϵ*^*w*^, corresponding to across-area connectivity being much more specialized than within-area, the dynamics in Equation 130 are dominated by across-area interactions. As shown earlier, directionality is in this case still led by the area receiving the strongest excitatory input (though it also acquire sensitivity to inhibitory inputs, see Supplementary Fig. 7b and 8). Conversely, when *ϵ*^*w*^≫ *ϵ*^*h*^, the dynamics in Equation 130 are shaped primarily by within-area interactions. Building on the analysis in Fig. 4h, if inhibitory inputs are comparable across areas, the area with stronger excitatory input is expected to lag behind the other. In this regime, differently tuned subpopulations within each area, being nearly decoupled, develop divergent dynamics. In particular, input-untuned subpopulations become largely locked to the activity of input-tuned subpopulations in the opposite area.

## Model simulations

All software was written in the Python programming language. Model circuit activity (Equation 3) was simulated using the Euler integration method. Below, we provide detailed information on the implementation and parameters used for the figures in the main text.

Figure 2. We simulated a network of *N* = 8 units with randomly generated, mean-zero Gaussian connectivity, and then arbitrarily assigned each neuron to one of two areas. The connectivity matrix was chosen to have only real eigenvalues. During the first epoch, the input vector associated with the active source was randomly generated. During the second epoch, the input vector was instead aligned with one of the eigenvectors of the connectivity matrix. During the third epoch, the input vector was aligned with one of the eigenvectors of the shuffled connectivity matrix. Each epoch lasted 30 time units. We simulated activity over 100 trials, each defined by a distinct initial condition and input realization. To estimate the cross-covariance function from simulated data, we first applied Equation 1 to every admissible pair of time indices *t* and *τ*, and then averaged the resulting values over *t*. The first 10 time units within each epoch, corresponding to highly non-stationary activity, were excluded. Panels C and D show the asymmetry score of cross-covariance functions computed between neurons in different areas.

Figure 3. We simulated a network of *N* = 4 units with randomly generated, mean-zero Gaussian connectivity. The connectivity matrix was chosen to only have real eigenvalues. During the first epoch, the input vector associated with the active source was taken as a linear combination of two eigenvectors of the connectivity matrix. During the second epoch, the input vector was instead aligned with one of the eigenvectors of the connectivity matrix. We simulated activity over 100 trials, with each epoch lasting 80 time units. The first 20 time units were excluded. Panels E-F refer to the same model circuit as in Fig. 2.

Figure 4. We considered a simple two-area excitatory–inhibitory network with parameters: *w* = 1.5, *h* = 0.5, *γ* = 0.33, and *k* = 1.01. In Fig. 4b, the input vector was parameterized as ***u*** = (1 + *δ*_1_*/*2, 1 + *δ*_2_*/*2, 1− *δ*_1_*/*2, 1 −*δ*_2_*/*2), where *δ*_1_ and *δ*_2_ correspond to the differences in input strength to the excitatory and inhibitory populations displayed in the horizontal and vertical axes, respectively. In Fig. 4c–d, the input vector corresponded to the configurations indicated by the magenta and violet diamonds in panel B. In Fig. 4e–f, we simulated 20.000 input configurations with components uniformly distributed between 0 and 1; configurations leading to the over-inhibited regime were excluded. In panel F, the inter-areal connectivity was set to *h*_FF_ = 1.3 and *h*_FB_ = 0.2 for the grey density plot, and *h*_FF_ = 0.2 and *h*_FB_ = 1.3 for the green one. Fig. 4g and H replicate the analysis presented in Fig. 4b, but with *k* = 1.2 (G) and *h* = 0 (H), respectively.

Figure 5. In Fig. 5a and b, we considered the same networks as in Fig. 4c and d. We simulated activity over 100 trials for 80 time units, and discarded activity within the first 20 time units. In Fig. 5c and d, the input vector was set to ***u*** = (1, *u*_I_, 0.3, 0), where *u*_I_ denotes the strength of the inhibitory input shown along the horizontal axis. Along the vertical axis, we vary the inter-areal coupling strength *h*. We plot activity statistics for the second area (qualitatively similar results apply to the first one).

Figure 6. We considered a two-area excitatory–inhibitory network with feature specialization. Parameters were identical to those in Fig. 4. We additionally set *ϕ* = 0.99, *ϵ*^*w*^ = 0.2, *ϵ*^*h*^ = 0.3, *u*_E1_ = 3.2, *u*_E2_ = 0.8, *u*_I1_ = *u*_I2_ = 1. In B-C and F-G, we simulated activity over 100 trials for 80 time units and discarded the first 20 time units. In panel G, we applied Partial Least Squares Regression to activity from the excitatory populations in the two areas, using the implementation from Python package sklearn with a single component. In D, we varied *ϵ*^*h*^ along the horizontal axis, and *ϕ* along the vertical one. We fixed *ϵ*^*w*^ = 0.99. In E and I, we varied *ϵ*^*h*^ along the horizontal axis, and *ϵ*^*w*^ along the vertical one. We fixed *ϕ* = 0.99. In H, the values of *u*_E1_, *u*_E2_, *u*_I1_ and *u*_I2_ were parametrized as in Fig. 4b; we fixed *ϕ* = 0.99.

## Analysis of V1-V2 simultaneous recordings

### Recordings

Animal procedures and recording details have been described in previous work [9, 13]. Briefly, neural activity was simultaneously recorded from populations in V1 (34 to 128 neurons) and V2 (12 to 84 neurons) from three anesthetized monkeys. The dataset analyzed here corresponds to a subset of the recordings in which the spatial overlap between the receptive fields of the V1 and V2 populations was optimized. Each trial started with a period of visual stimulation during which monkeys were presented with drifting sinusoidal gratings (total of eight equally spaced orientations), followed by a period without stimulation. The duration of the evoked and spontaneous periods was 1280 ms and 1500 ms, respectively. Every recording session included 400 repetitions of each stimulus, yielding a total of 3200 trials. The dataset includes five recording sessions.

## Data analysis

We treated data corresponding to stimuli of different orientations as different datasets (total of 40). We excluded neurons that fired less than 0.5 spikes/s on average across all trials. Spiking activity was centered by subtracting each neuron’s trial-averaged activity over time, as well as its time-averaged activity on every trial.

Feature-unspecific axes for V1 and V2 populations were defined as normalized vectors with identical weights for all neurons in both populations. To define feature-specific axis, we first computed each neuron’s orientation tuning curve *F*_*i*_(*θ*), defined as the average spike count during the stimulus presentation. We then computed an orientation selectivity index (OSI) for every neuron *i* in V1 and V2, defined as:

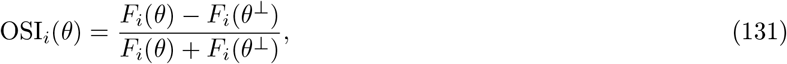

where *θ* and *θ*^*⊥*^ denote the presented stimulus orientation and its orthogonal counterpart, respectively. Feature-specific axes for V1 and V2 were obtained by taking the corresponding population vectors of OSIs, subtracting their means, and normalizing them. Using this definition, neurons with positive weights (resp. negative) form the population tuned to stimulus *θ* (resp. *θ*^*⊥*^). Alternatively, feature-specific axes can also be obtained by setting opposite-sign weights for two populations in each area, with the boundary defined as the sign or median of the population OSIs. The latter approach naturally yields feature-specific axes that are orthogonal to the feature-unspecific ones. These alternatives produce axes closely related to those used in our model (Fig. 6) and lead to qualitatively similar results to those shown in Fig. 7. To compute unspecific and specific latents in both areas, we projected the V1 and V2 residual activity onto the corresponding axes.

Following [13], cross-covariance functions were obtained in a time-resolved way, by considering 80 ms windows advanced in steps of 40 ms, and then applying Equation 1 for a range of lag values. Within each window, activity was binned with non-overlapping bins of 2 ms. The dominant cross-covariance functions were obtained using a similar scheme, and by re-running the PLS algorithm for every combination of time and lags. All results for the two epochs display averages performed over the time index within specific windows in which activity was close to stationary: [120, 360] ms for the evoked epoch, which includes the early evoked period analyzed by [13], and [1960, 2440] ms for the spontaneous epoch. To capture inter-areal interactions occurring at fast timescales, which dominate directionality in the early stimulus epoch, the asymmetry scores were computed in the lag windows [-8, 8] ms. Our results are robust to other windows choices, over a reasonable range.

A potential concern was that the non-negativity of activity in the spiking data (but not in the models) could artificially bias feature-unspecific components to have larger amplitude than feature-specific ones. As a consequence, the axes extracted by PLS could also appear more aligned with the feature-unspecific direction simply because these components carry larger variance. To address this concern, we preprocessed the data by centering activity over time and trials (see above). These pre-processing steps did not alter the qualitative outcome of the analysis. In addition, we repeated the analysis using canonical correlation analysis (CCA) instead of PLS. CCA is designed to capture shared co-modulations across areas independently of the absolute amplitude of the latent signals, and was in fact originally used in [13]. This alternative analysis yielded qualitatively similar results.

## Data availability

The V1–V2 data used in this study are available in the CRCNS data sharing website, at https://doi.org/10.6080/K0B27SHN.

## Code availability

Implementations of all simulations and algorithms used in this study is available at https://github.com/fmastrogiuseppe/InputDrivenDirectionality.

## Acknowledgments

The authors would like to thank Matthijs oude Lohuis and Alfonso Renart for discussion and feedback on the manuscript.

## Author Contributions Statement

F.M. and J.C. performed the formal and computational analyses, analyzed the neural recordings data, and wrote the manuscript. F.M., B.M.Y., A.K. and C.K.M. supervised the work. B.M.Y., A.K. and C.K.M. provided feedback on the manuscript.

## Competiting Interests Statement

The authors declare no competing interests.

## Supplementary Information for

**Supplementary Fig. 1:**
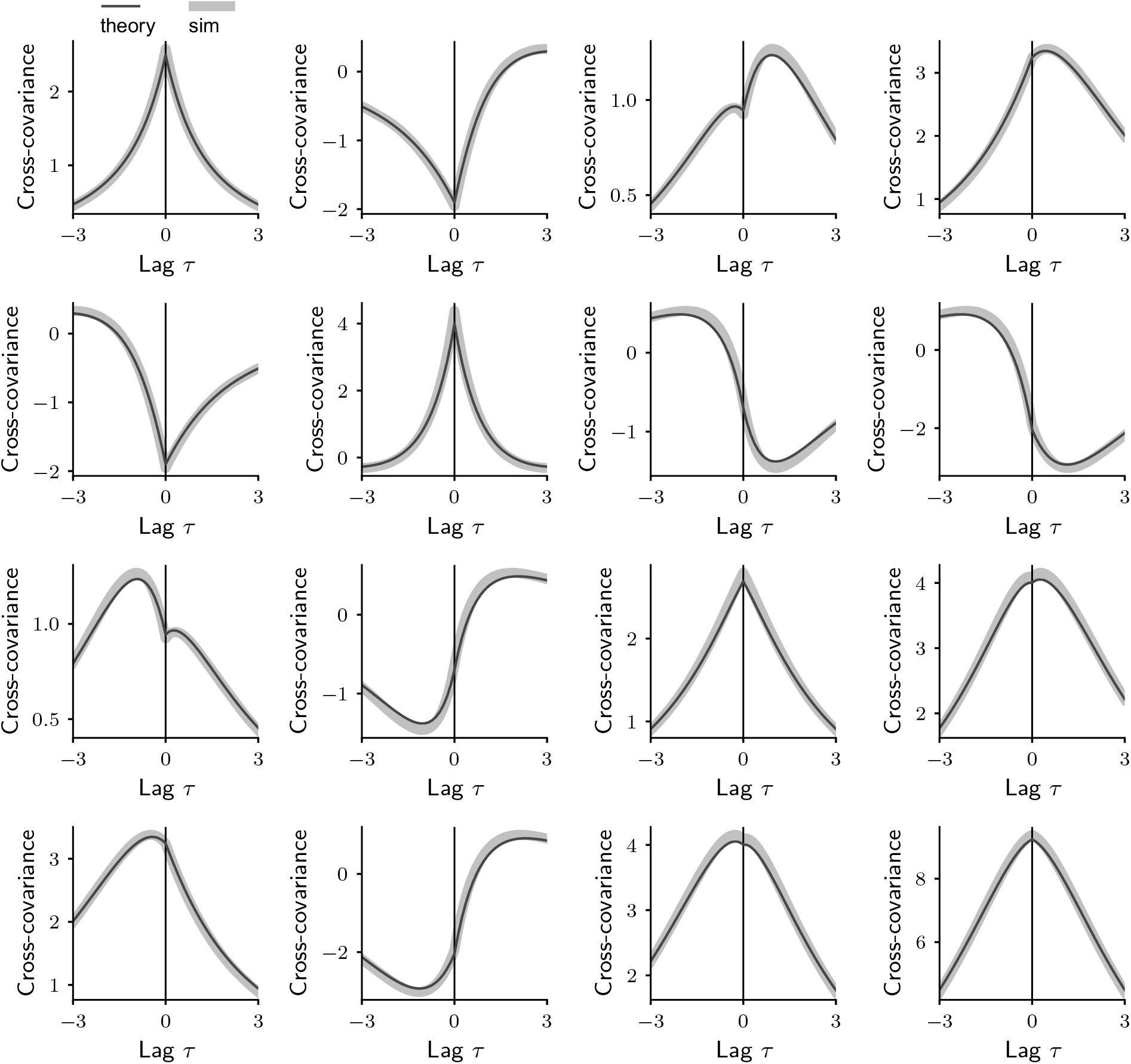
Full set of covariance functions for a representative circuit with *N* = 4 units. Both recurrent and input connectivity (Eq. 3) are generated at random. The panel in the *i*-th row and *j*-th column shows the cross-covariance between units *i* and *j* (Eq. 1). Diagonal panels display single-unit auto-correlations, exhibiting different effective timescales. Off-diagonal panels display cross-covariances, which are temporally asymmetric. Thick light lines indicate covariances computed from simulated activity, while thin dark lines show covariances computed from theoretical expressions (Eqs. 42 and 43). The two are in good agreement.

**Supplementary Fig. 2:**
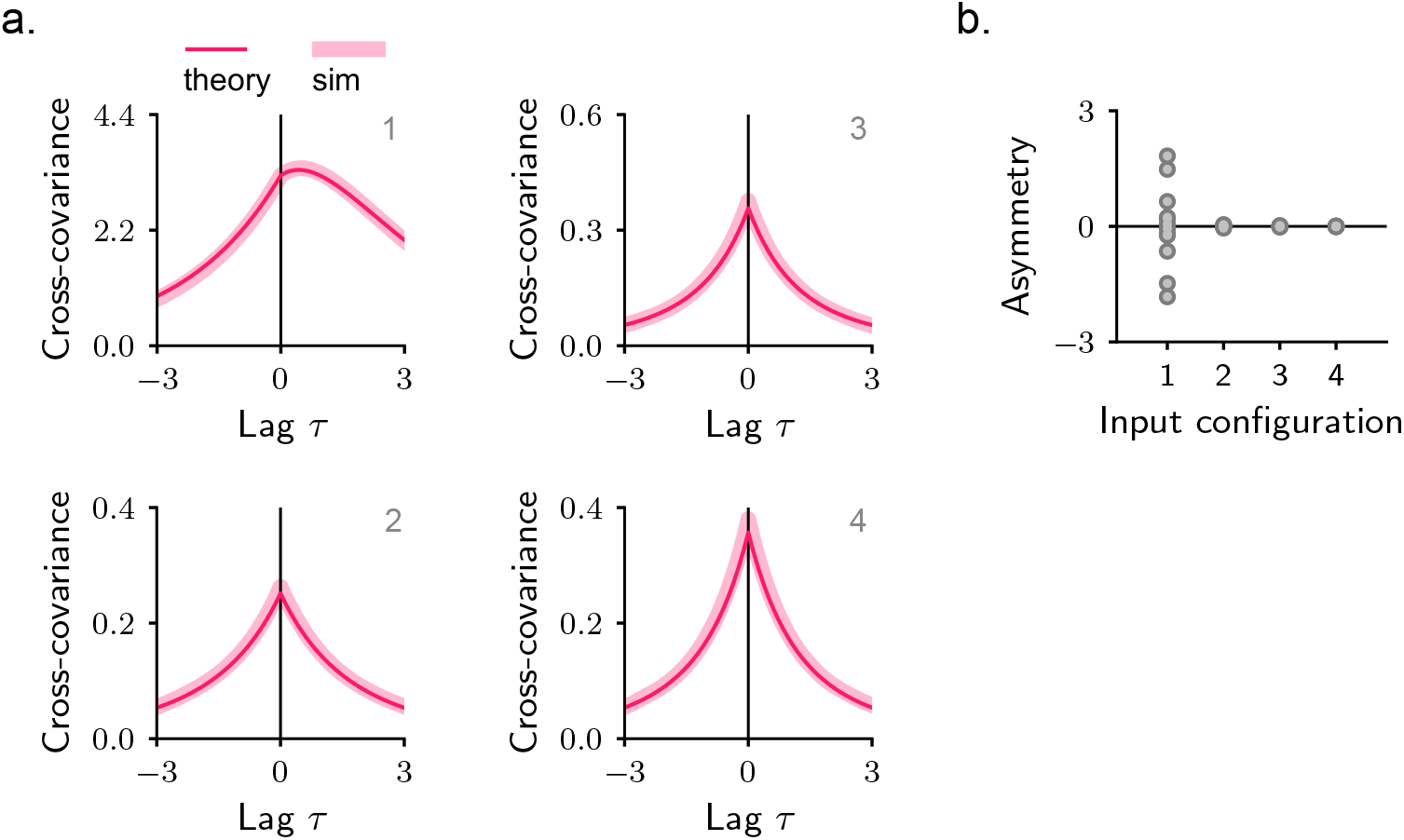
Directionality in circuit models with high-dimensional inputs. **a**. Example cross-covariances from four different circuits, all sharing the same randomly generated recurrent connectivity, but differing in their input connectivity. In panel 1, *U* is generated randomly. In panel 2, *U* = *P*, where *P* contains the eigenvectors of the recurrent connectivity as columns. In panel 3, *U* = *PD*^1*/*2^, where *D* is a diagonal matrix with random entries along the diagonal. Finally, in panel 4, *U* = *PD*^1*/*2^*R*, where *R* is an orthogonal matrix. For input configurations 2, 3, and 4, the theory predicts symmetric cross-covariances across all unit pairs (see Methods). Thick light magenta lines indicate cross-covariances computed from simulated activity, while thin dark magenta lines show cross-covariances computed from theoretical expressions (Eqs. 42 and 43); the two are in good agreement. **b**. Directionality, quantified via the asymmetry score, for all cross-covariances in the simulated circuits. The asymmetry score remains close to zero for all cross-covariances in input configurations 2, 3, and 4.

**Supplementary Fig. 3:**
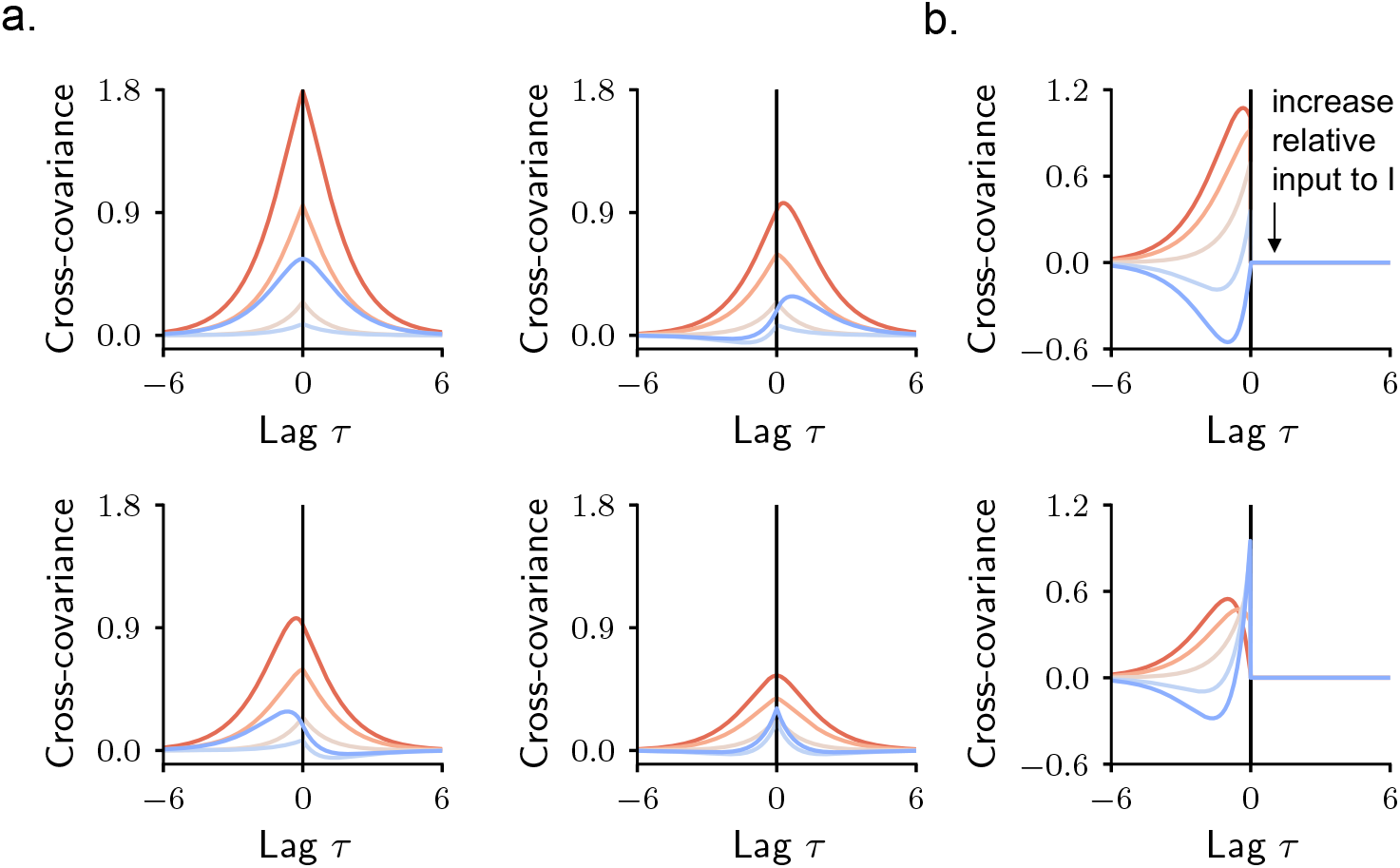
Directionality in single-area excitatory–inhibitory models. We consider a single-area network as described in Methods, with local connectivity parameters as in Fig. 4. Different lines correspond to different input configurations, with red (resp. blue) shades indicating stronger input to the E (resp. I) population. The input vector is constructed as ***u*** = (cos(*θ*), sin(*θ*)), where *θ* ranges from 0 (dark red) to 90 degrees (dark blue). **a**. Full set of covariances. Diagonal panels show the auto-covariance of E (top) and I populations (bottom); off-diagonal panels show their cross-covariance. Values are computed from theoretical expressions (Eqs. 42, 43). **b**. Activity–input cross-covariance (Eq. 63) for E (top) and I (bottom) populations. Values are computed from theoretical expressions (Eq. 65). Note that when the input to the E population is stronger, activity exhibits large amplitude, and both E and I populations co-fluctuate with the external input. In contrast, when the input to the I population dominates, activity has smaller amplitude, and fluctuations in the E population are anti-correlated with the external input. We refer to such configurations as the over-inhibited regime.

**Supplementary Fig. 4:**
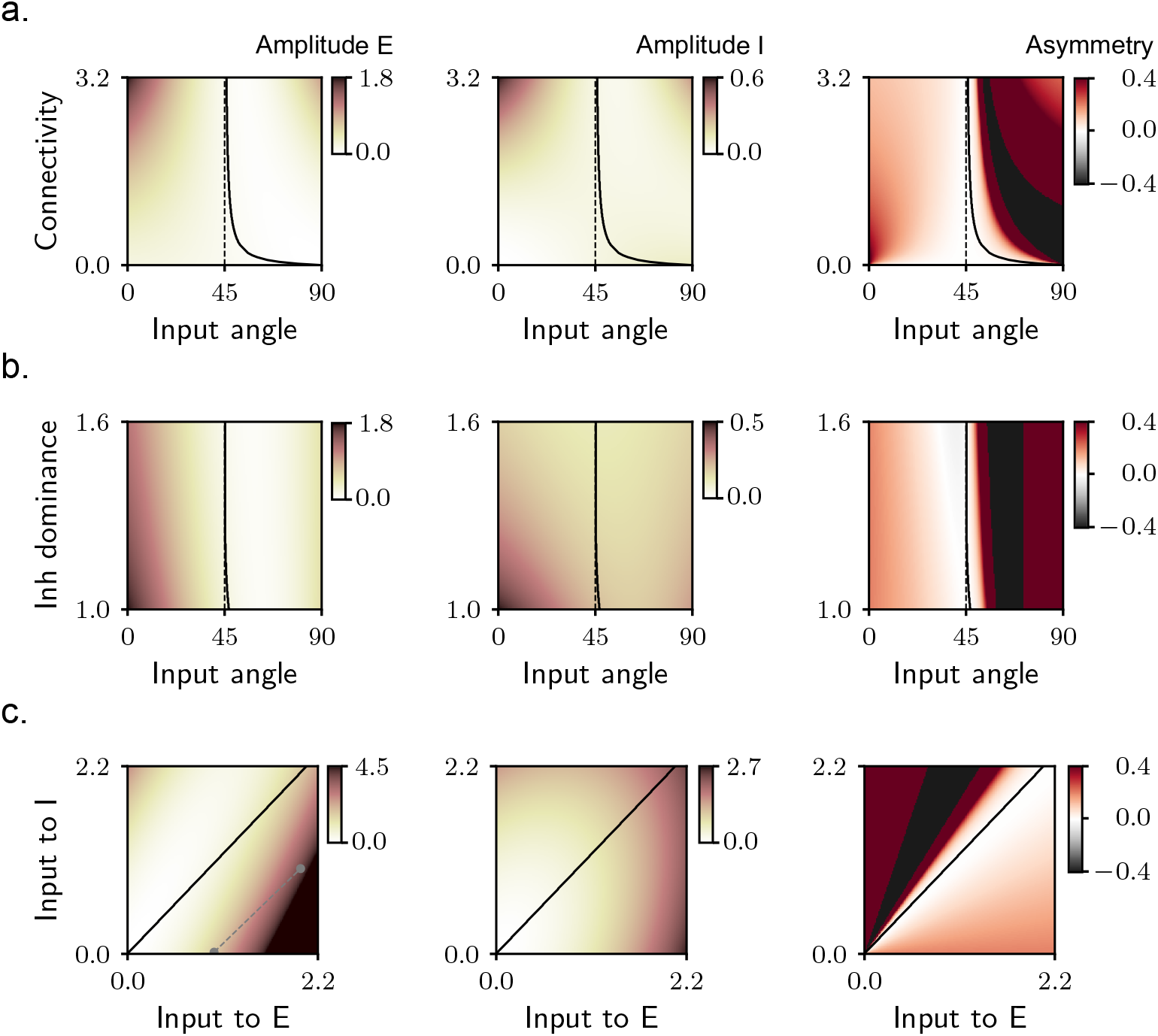
Systematic characterization of directionality in single-area excitatory–inhibitory models. We consider a single-area network, with local connectivity parameters as in Fig. 4. Left column: amplitude of activity in the excitatory (E) population, measured as variance of activity fluctuations. Center column: amplitude of activity in the inhibitory (I) population. Right column: temporal asymmetry of the cross-covariance function between E and I populations, quantified by the asymmetry score. All values are computed from theoretical expressions (Eqs. 42, 43). **a**. Activity statistics as a function of the input direction and the recurrent connectivity strength, *w*. The input vector is defined as ***u*** = (cos(*θ*), sin(*θ*)), with *θ* ranging from 0 to 90 degrees. The dashed black line marks the value of *θ* at which E and I populations receive identical inputs; in this case, the theory predicts vanishing asymmetry. For parameter values to the right of the solid black line, activity lies in the over-inhibited regime, for which cross-covariances are strongly asymmetric. **b**. Activity statistics as a function of the input vector angle, *θ*, and the relative inhibition dominance, *k*. Details as in a. **c**. Activity statistics as a function of the input strength to E and I populations. For parameter values to the left of the solid black line, activity lies in the over-inhibited regime. Grey dots in the left column refer to the small-*h* mathematical analysis in Methods. Moving from one dot to the other, both E and I inputs increase by the same amount. Nevertheless, activity amplitude is substantially larger for the top-right dot, corresponding to stronger E inputs.

**Supplementary Fig. 5:**
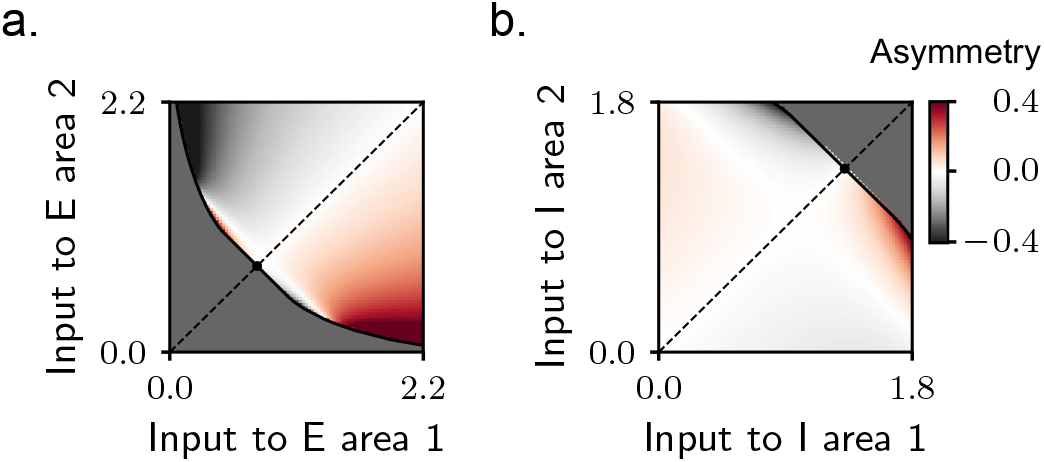
Further characterization of across-area directionality in two-area excitatory–inhibitory models. We consider a simple two-area network as in Fig. 4. **a**. Cross-covariance asymmetry score as a function of the input strength to the excitatory populations in area 1 (horizontal axis) and area 2 (vertical axis). The input to inhibitory populations is fixed at 1. The dashed black line marks input configurations that are symmetric across the two areas, for which directionality necessarily vanishes. The dark grey region corresponds to the regime in which strong inhibitory inputs suppress excitatory activity. The tiny black dot indicates the theoretical prediction for the boundary of this region (see Methods). Input configurations far from the diagonal consistently exhibit large asymmetry values, confirming that differences in excitatory inputs strongly modulate directionality (see also Fig. 4b). **b**. Same as a, but for inputs to the two inhibitory populations. The input to excitatory populations is fixed at 1. Here, configurations away from the diagonal display small asymmetry scores, indicating that differences in inhibitory drive weakly affect directionality (Fig. 4b).

**Supplementary Fig. 6:**
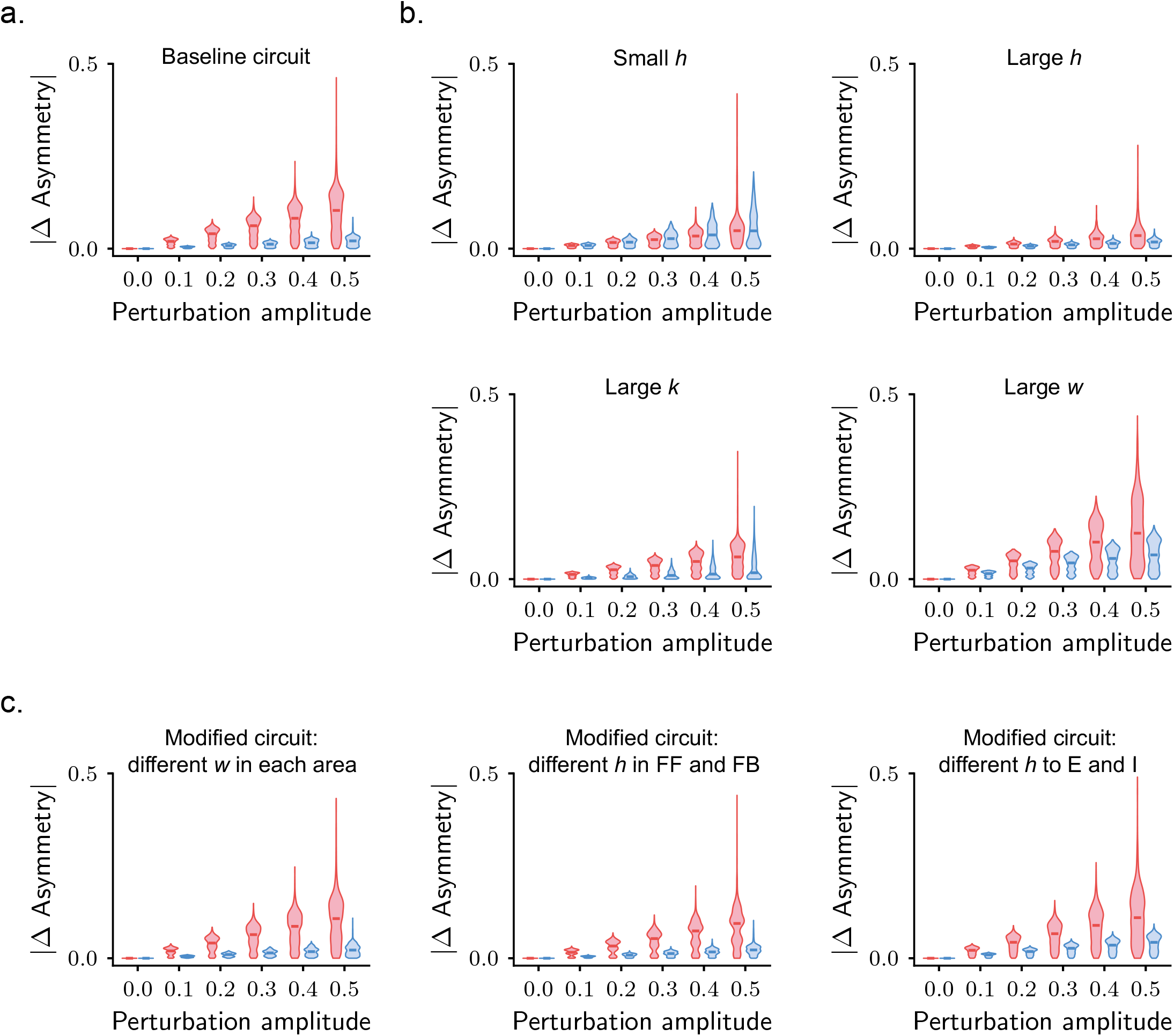
Differential sensitivity of directionality to inputs targeting excitatory and inhibitory populations: robustness and dependence on network parameters. We consider a simple two-area network as in Fig. 4. To systematically investigate how the results in Fig. 4 depend on network parameters, we quantify the differential sensitivity of directionality in a different form. Starting from a random input configuration, we investigate the change in directionality obtained following a perturbation to the input strengths into the excitatory versus the inhibitory populations. The initial input vector is generated from a uniform distribution, with range [0.5, 1]. To each initial vector, we apply a set of additive perturbations to excitatory and inhibitory populations. The input perturbations are defined as ***δ***_E_ = (*δ* cos *θ*, 0, *δ* sin *θ*, 0) and ***δ***_I_ = (0, *δ* cos *θ*, 0, *δ* sin *θ*), where *δ* and *θ* take equally spaced values in the ranges [0, 0.5] and [0^*°*^, 360^*°*^], respectively. We then compute the cross-covariances obtained for the unperturbed and perturbed input vectors and the corresponding difference in asymmetry scores. Violin plots show the distributions of absolute differences in asymmetry score, |ΔAsymmetry |, for input perturbations to the excitatory (red) and inhibitory (blue) populations, as a function of perturbation amplitude. Input vectors leading to the over-inhibited regime are excluded, and each violin contains 2000 points. Directionality is disparately sensitive to differences in inputs targeting the excitatory and inhibitory populations when the red and blue distributions (in particular, their medians, indicated by the red and blue dashes) are different. In `a-c`, we focus on the simplest two-area circuit model described in Methods, and consider different sets of network parameters, for which the differential sensitivity is expected to be stronger or weaker. In `c`, we consider circuit configurations in which we relax some of the assumptions of the simple model, as described in Methods, and test whether the differential sensitivity also extends to these models. **a**. Network parameters as in Fig. 4. The circuit is characterized by sufficiently strong across-area connectivity (*h* = 0.5) with respect to within-area one (*w* = 1.5) and a close balance between excitation and inhibition (*k* = 1.01). **b**. We start from the baseline circuit in a, and then set: weaker across-area connectivity (*h* = 0.25, top left); stronger across-area connectivity (*h* = 0.95, top right); stronger imbalance between the local strength of excitation and inhibition (*k* = 1.25, bottom left); stronger within-area recurrence (*w* = 5, bottom right). For all these parameter settings, differential sensitivity is weaker than in a, as predicted by the theoretical analysis. **c**. Circuit models where the simplifying connectivity assumptions are relaxed (see Methods). Left: different strength of within-area excitatory connections in the two areas (*w*_1_ = 1 and *w*_2_ = 2). Center: different across-area connectivity in the feedforward and feedback directions (*h*_FF_ = 1.3 and *h*_FB_ = 0.2, respectively). Right: different strength of across-area connections to the excitatory and inhibitory populations, given by *h* and *αh*, respectively (*h* = 0.6, *α* = 0.9). All other network parameters are as in Fig. 4. The differential sensitivity is robustly observed across these modified circuit models, suggesting that it is an emergent property of multi-area circuits with locally balanced excitation and inhibition and sufficiently large excitatory across-area connectivity.

**Supplementary Fig. 7:**
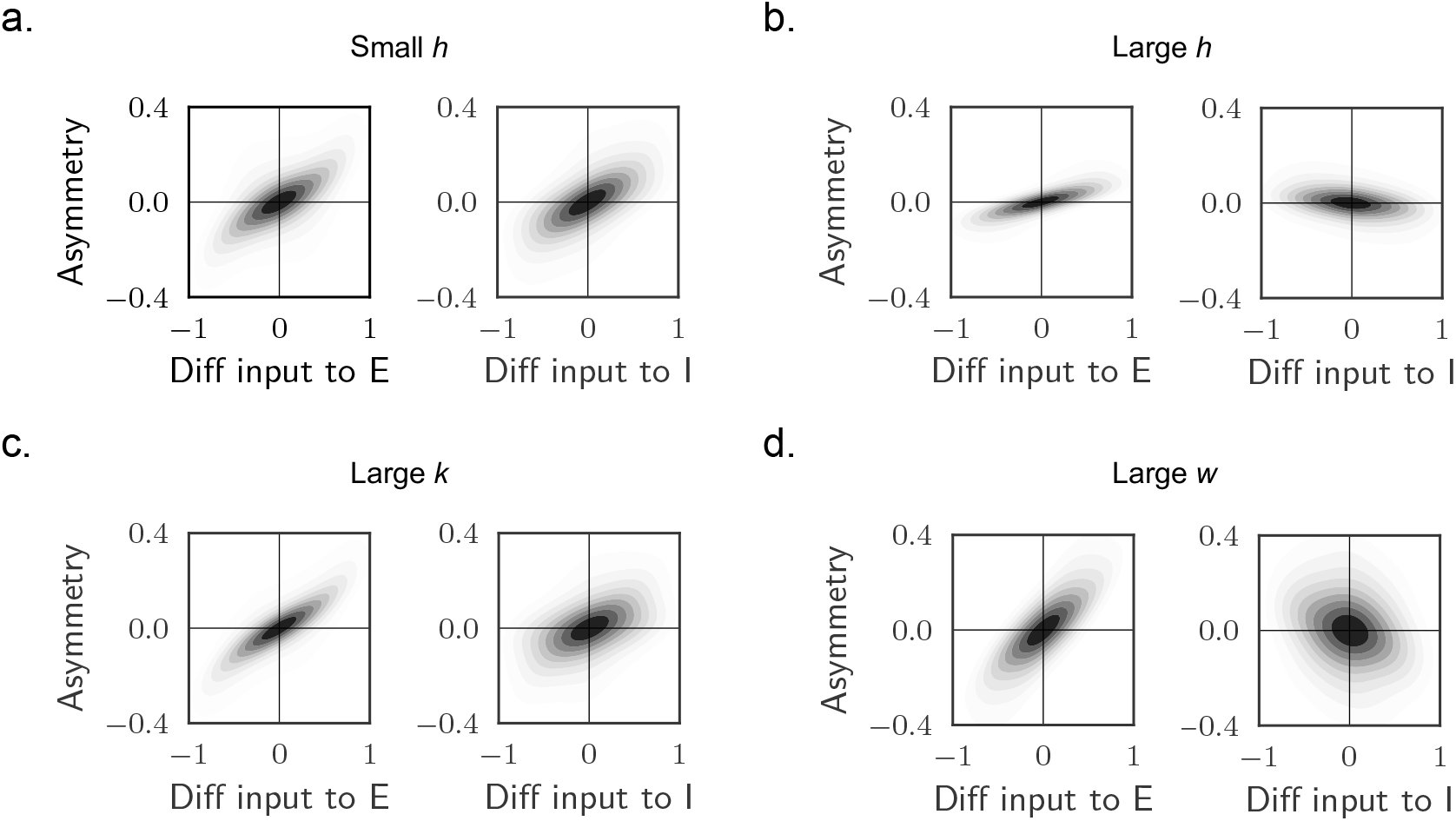
Influence of network parameters on directionality sensitivity to inputs targeting excitatory and inhibitory populations. We consider a simple two-area network as in Fig. 4. Fig. 4e-f shows that there exists a monotonic relationship between the difference in input to the excitatory populations and the cross-covariance asymmetry score, which is abolished for differences in input to the inhibitory populations. To assess how this result depends on network parameters, we repeat this analysis for different sets of network parameters. Density plots show the asymmetry scores obtained for a set of 20000 random input configurations, with components uniformly distributed between 0 and 1, as a function of the difference in input to the excitatory (left) and inhibitory (right) populations. Input vectors leading to the over-inhibited regime were excluded. We start from the baseline parameters used in Fig. 4 and then set: **a**. weaker across-area connectivity (*h* = 0.25); **b**. stronger across-area connectivity (*h* = 0.95); **c**. stronger imbalance between the local strength of excitation and inhibition (*k* = 1.25); **d**. stronger within-area recurrence (*w* = 5). Across all these parameter settings, directionality exhibits sensitivity to differences in inputs to inhibitory populations – although weaker than for excitatory ones, see also Supp. Fig. 6 – consistent with our theoretical predictions.

**Supplementary Fig. 8:**
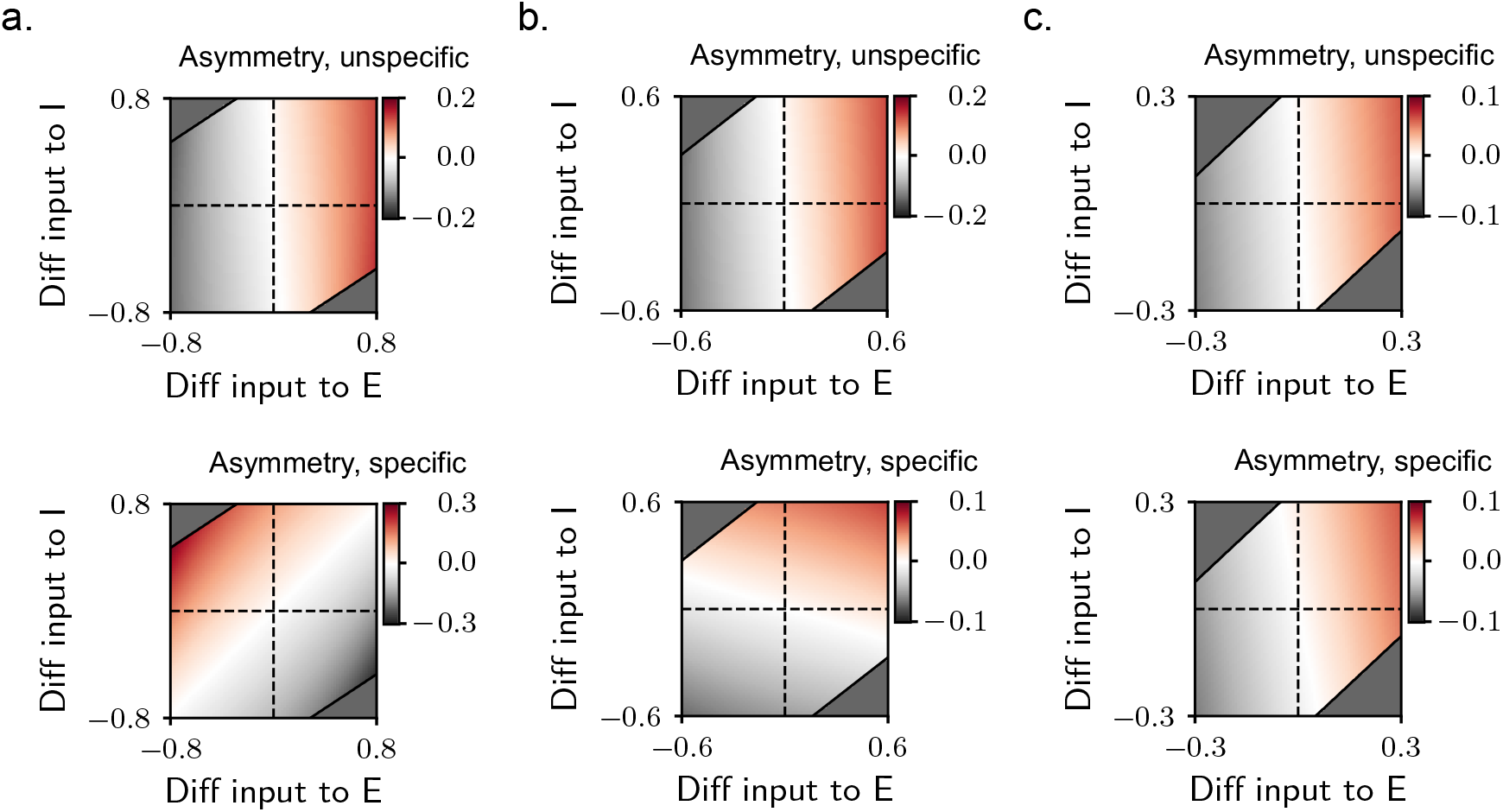
Across-area directionality for feature-unspecific and specific latent signals. We consider a feature-specific two-area network as in Fig. 6. We parametrize input configurations as in Fig. 4b, while fixing input specificity to *ϕ* = 0.99. Directionality is quantified as the asymmetry score of the cross-covariance function computed from feature-unspecific latents (top) and feature-specific ones (bottom). **a**. Circuit with very low feature specialization in inter-areal connectivity: *ϵ*^*h*^ = 0.02. **b**. Circuit with intermediate feature specialization: *ϵ*^*h*^ = 0.3. **c**. Circuit with high feature specialization: *ϵ*^*h*^ = 0.7. In all panels, within-area specialization is fixed at *ϵ*^*w*^ = 0.2. The qualitative behaviour of directionality for feature-unspecific (resp. specific) latents is independent (resp. depenendent) on changes in inter-areal connectivity.

